# Cell type-specific immune regulation under symbiosis in a facultatively symbiotic coral

**DOI:** 10.1101/2024.06.20.599951

**Authors:** Maria Valadez-Ingersoll, Hanny E. Rivera, JK Da-Anoy, Matthew R. Kanke, Kelly Gomez-Campo, M. Isabel Martinez-Rugerio, Julian Kwan, Ryan Hekman, Andrew Emili, Thomas D. Gilmore, Sarah W. Davies

**Affiliations:** Boston University, Department of Biology; Boston, MA, US; Gingko Bioworks, Boston, MA, USA; Amgen Research, Research Bioimics, South San Francisco, CA, USA; Pennsylvania State University, Department of Biology, State College, PA, USA; Helmholtz Institute for Functional Marine Biodiversity (HIFMB), Ammerländer Heerstraße 231, 26129 Oldenburg, Germany; Department of Biochemistry, Boston University Chobanian & Avedisian School of Medicine, Boston, MA, USA; Division of Oncological Sciences, Knight Cancer Institute, Oregon Health & Science University, Portland, OR, USA

**Keywords:** coral, symbiosis, immunity, single-cell RNA-sequencing, proteomics

## Abstract

Many cnidarians host single-celled algae within gastrodermal cells, yielding a mutually beneficial exchange of nutrients between host and symbiont, and dysbiosis can lead to host mortality. Previous research has uncovered symbiosis tradeoffs, including suppression of immune pathways in cnidarians hosting intracellular algae and correlations between symbiotic state and pathogen susceptibility. Here, we used a multiomic approach to characterize symbiotic states of the facultatively symbiotic coral *Oculina arbuscula* by generating genotype-controlled fragments of symbiotic and aposymbiotic tissue. 16S metabarcoding showed no difference in bacterial communities between symbiotic states. Whole-organism proteomics revealed differential abundance of proteins related to immunity, confirming immune suppression during symbiosis. Finally, single-cell RNAseq identified diverse cell clusters within seven cell types across symbiotic states. Specifically, the gastrodermal cell clusters containing algal-hosting cells from symbiotic tissue had higher expression of nitrogen cycling and sugar transport genes than aposymbiotic gastrodermal cells. Furthermore, differential enrichment of immune system gene pathways and lower expression of genes involved in immune regulation were observed in these gastrodermal cells from symbiotic tissue. However, no differences in immune gene expression in the immune cell cluster were observed between symbiotic states. This work reveals a compartmentalization of immune system regulation in specific gastrodermal cells in symbiosis, which may limit symbiosis tradeoffs by simultaneously dampening immunity in algal-hosting cells while maintaining general organismal immunity.

## INTRODUCTION

Symbioses exist across the Tree of Life along a spectrum from parasitisms – in which one symbiotic partner benefits at the cost of the other partner – to mutualisms – in which both partners benefit from the relationship (1). Endosymbioses, symbiotic relationships where one organism resides within another, pose immunity challenges to hosts as they must differentiate between self, mutualistic symbionts, and pathogens (2,3). In mutualisms, the host immune system is often regulated in order to allow for the existence of the endosymbiont (4), whereas in parasitisms the endosymbiont can evade detection by the host’s immune system (5). Additionally, in mutualisms, hosts must tightly regulate the density of their symbionts in order to prevent overcrowding, which can lead to a shift from mutualism to parasitism (6).

Reef-building corals participate in a mutually beneficial symbiosis with algae that reside inside their gastrodermal cells, wherein the algae carry out photosynthesis and translocate fixed carbon sugars to the host (7,8). In return, the host provides CO_2_, as well as nitrogen and phosphorus species to the symbiont (9,10). Translocation of nitrogen from the host to the symbiont has been proposed as a mechanism to promote nutrient exchange while controlling algal proliferation via nitrogen limitation to prevent overcrowding (11). As symbiont densities increase, the greater nutrient requirements of the algae lead to resource competition with the host, thereby destabilizing the mutualism (12,13). In support of this model, whole-organism RNA-sequencing has revealed differences in the expression of genes involved in sugar transport and nitrogen cycling across cnidarians in and out of symbiosis (14), and enrichment of nitrogen deprivation genes in algae exhibiting greater cell densities when associated with the sea anemone *Exaiptasia pallida* (11). These whole organism studies reflect the importance of carbon and nitrogen transport between symbiotic partners in the maintenance of this symbiosis.

In stony corals, environmental change can lead to loss of the algal symbiont in a process termed coral bleaching (15). If prolonged, this dysbiosis can lead to host mortality. Since this symbiotic relationship in corals is often obligate, it is challenging to disentangle the effects of dysbiosis from nutritional losses (16). Accordingly, facultatively symbiotic cnidarian models, which are viable and naturally occur in both symbiotic and aposymbiotic states, have been useful for understanding mechanisms that promote symbiosis establishment, maintenance, and loss, from both nutritional and immune system perspectives (14,16–18). In contrast to obligate symbioses, facultatively symbiotic hosts can exist in an aposymbiotic state (with few or no algal symbionts) and can buffer nutrient loss through heterotrophy. This facultative nature allows for studies of the influence of symbiosis on host molecular function in the absence of starvation stress experienced by bleached obligate corals (19). Importantly, the aposymbiotic state can be induced in facultatively symbiotic cnidarians in the laboratory through chemical (with menthol) and heat treatments, additionally allowing for the ability to control for genetic background. The sea anemone *E. pallida* is a facultatively symbiotic cnidarian that has served as a useful model for understanding how the host immune system is modulated in the presence of the algal symbiont (20,21). Additionally, facultatively symbiotic stony corals, such as *O. arbuscula* (14) and *Astrangia poculata* (22), are emerging models for studying these processes in calcifying cnidarians.

Research using these facultatively symbiotic models has identified pathways within the host immune system as key for the cnidarian-algal symbiosis. For example, activation of the transcription factor NF-κB pathway, which has been implicated in immunity across metazoans, has been shown to be downregulated in symbiotic compared to aposymbiotic hosts (14,20). Additional evidence suggests that the attenuation of immune pathways associated with symbiosis corresponds to reduced organismal immunity (23). However, this dampening of immunity is not always correlated with increased susceptibility to pathogen challenge (24,25). For example, recent work has shown that the NF-κB pathway can be independently modulated by cnidarians capable of symbiosis under energetic limitation (26). Additionally, many antioxidant and oxidative response pathways are constitutively higher in symbiotic compared to aposymbiotic cnidarians (27). Taken together, a nuanced picture of the symbiotic cnidarian immune system has emerged, with the question remaining of how the immune system challenge of endosymbiosis is mitigated without compromising whole-organism immunity.

Herein, we have used a multiomic approach to investigate symbiosis-immunity tradeoffs in the facultatively symbiotic coral *O. arbuscula*. Whole-organism proteomics confirms broad immune dampening in symbiosis. Single-cell RNA-seq of genotype-controlled fragments was used to profile *O. arbuscula* cellular diversity under different symbiotic states. We provide evidence for how tradeoffs of endosymbiosis and immunity manifest by gene expression rewiring across different cell states. Strikingly, we found gene expression differences in immune pathways occur in gastrodermal cells capable of hosting symbiotic algae. These results suggest that cell type-specific dampening of immunity explains how some symbiotic organisms balance the immune tradeoffs of symbiosis with organismal immunity.

## MATERIALS AND METHODS

Detailed information on sample preparations, bioinformatic analyses, and R packages can be found in the **Supplemental Materials**.

### Coral husbandry and manipulation of symbiotic state

Symbiotic (in symbiosis with *Brevioulum psygmophilum*) (28) colonies from seven genetic backgrounds (genets A-G) of *Oculina arbuscula* were collected at Radio Island Jetty, North Carolina and have been maintained at Boston University (BU) in common garden aquaria since May, 2018. Aposymbiotic branches were generated via menthol bleaching. Aposymbiotic status was confirmed by a lack of symbiont autofluorescence under fluorescence microscopy (Leica M165 FC), and aposymbiotic branches were transferred back to common garden aquaria and maintained for at least 2 months of recovery prior to physiological and multiomic profiling.

### Tissue removal for symbiont cell quantification

To measure symbiont cell densities in symbiotic and aposymbiotic *O. arbuscula*, small branches were fragmented from symbiotic (N=10) and aposymbiotic (N=10) samples from five genets. Tissue was removed via airbrushing, homogenized, and symbiont counts were normalized to skeletal surface area using an Einscan-SE scanner. Statistical differences between symbiotic and aposymbiotic fragments were calculated using the Kruskal-Wallis rank sum test, as assumptions of normality were not met.

### Coral spectroscopic determinations

Light absorption capacities of symbiotic and aposymbiotic *O. arbuscula* were compared using reflectance and absorptance (29–31). Coral reflectance (R), the fraction of light reflected, was measured between 400-750 nm in intact fragments using a miniature spectroradiometer (Flame-T-UV-Vis, Ocean Optics Inc.). Reflectance was expressed as the ratio of the measurement from the tissue surface relative to the reflectance of a bleached *O. arbuscula* skeleton. Coral absorptance (A), which describes the fraction of incident light absorbed by coral tissue, was calculated from the reflectance spectra as A = 1 – R (29,31). The absorptance peak of chlorophyll a (Chl a) at 675 nm was calculated as A_675_ = 1 – R_675_, assuming that transmission through the skeleton is negligible.

### Bacterial community profiling

To identify bacterial communities associated with symbiotic and aposymbiotic fragments, one symbiotic and one aposymbiotic fragment from five genotypes were flash frozen, and a subsample was preserved in ethanol for 16S metabarcoding (N=10). Metabarcoding libraries were generated using a series of PCR amplifications for the V4/V5 region of the bacterial 16S rRNA gene (32,33). Five negative controls were prepared and used to remove contamination. Samples were sequenced on an Illumina Miseq at Tufts University Core Facility (paired-end 250 bp).

16S read processing inferred 661 amplicon sequence variants (ASVs) with assigned taxonomy across samples. Bacterial communities across symbiotic states were compared via alpha diversity (Shannon index, Simpson’s index, ASV richness, and evenness) and beta diversity (PCoA with Bray–Curtis dissimilarities). Alpha diversity metrics were compared using linear mixed effects models with symbiotic state as the predictor and a random effect of genotype. The effect of symbiotic state on beta diversity was assessed using the function *betadisper* (vegan package; v2.6-4 (34)). DESeq2 v1.40.2 (35) explored differentially abundant ASVs to ensure that subtle differences in specific taxa were not overlooked.

### Proteomic profiling

Mass spectroscopy (MS) was used to identify differentially enriched proteins from total protein isolated from fragments of four symbiotic genets and three aposymbiotic genets. Total protein was isolated as described previously (36) and was stored at −80 ℃ before analysis by MS.

For MS, tryptic peptide mixtures were analyzed by nano-scale high-performance liquid chromatography (Proxeon EASY-Nano system, Thermo Fisher Scientific) coupled with online nanoelectrospray ionization tandem MS (Q-Exactive HF-X mass spectrometer; Thermo Fisher Scientific) (37). For protein identification and analysis, data files were searched using the workflow of MaxQuant version 2.4 (http://www.maxquant.org/) under standard settings using the *Astrangia poculata* genome (22). A false discovery rate (FDR) threshold of 1% was used to filter candidate peptides and protein identifications. Following data filtration, normalization, and clustering with Omics Notebook (38), 2,191 proteins were retained. Differential abundance analysis of proteomic profiles between aposymbiotic and symbiotic samples was based on a moderated t-test (38) in which proteins were considered differentially abundant if they had a Bonferroni adjusted *p*-value < 0.1. Raw intensity counts were *rlog-*normalized in DESeq2 (35) and visualized using a Principal Component Analysis (PCA) using vegan v2.6-4 (34). Effects of symbiotic state and genet were assessed using a PERMANOVA with the adonis2 function in vegan v2.6-4 (34). Predicted *O. arbuscula* peptides were searched against the human proteome v.11.5 from the STRING v.11 database (39) with an e-value cut-off of 1×10^-5^. Protein–protein interactions of select differentially expressed proteins (FDR < 0.1) were retrieved from the STRING v.11 database (39). Interaction networks were visualized using Cytoscape v.3.7.2 (40).

### Single-cell RNA sequencing

To create single-cell libraries, live cells from one symbiotic and one aposymbiotic branch of genotype F of *O. arbuscula* were sampled for single-cell isolation and 10X cDNA sequencing (41). A detailed protocol for cell isolation can be found on protocols.io at DOI: dx.doi.org/10.17504/protocols.io.rm7vzkx72vx1/v1 and is also described in the **Supplemental Materials**. To minimize cell death, cells were not sorted prior to processing. Cell isolation samples were analyzed by BU’s Single Cell Sequencing Core Facility. The symbiotic sample had a concentration of 2,625 cells/μl and a viability of 81.6%. The aposymbiotic sample had a concentration of 3,313 cells/μl and a viability of 86.4%. These values are largely in line with other single cell preparation methods reported in the literature (e.g., viability threshold of 80%, *Nematostella vectensis* (42,43)).

Libraries were generated following the 10X Genomics Chromium Single Cell 3’v3 protocol, and quality was assessed via a Bioanalyzer. Samples were pooled in equimolar concentration and sequenced on an Illumina NextSeq 2000 (P3 100 kit) with a target of 50,000 reads per cell. CellRanger (v7.2.0 (44)) processed reads, which were aligned to concatenated genomes of *Astrangia poculata* (host) (22) and the algal symbiont *Breviolum psygmophilum* (14). Reads aligning confidently to the host and symbiont references were used to generate CellRanger output files. Default CellRanger cell calling algorithms were employed. The *A. poculata* genome was used as a reference for mapping because there is currently no published genome for *O. arbuscula,* and because alignment rates to a *de novo* assembled transcriptome from the 10X data generated here and to a previously published *O. arbuscula* transcriptome were of lower quality (**Supplementary Dataset 1**) (14,19,45). In addition, it is worth noting that species in the genera *Astrangia* and *Oculina* are closely related phylogenetically (19), and previous work has found that the *O. arbuscula* population from Radio Island, NC (source for this study) is more genetically similar to the *A. poculata* population from Woods Hole, MA (coral genome source) than it is to other, more southern *Oculina* populations (45).

Following alignment, host reads from symbiotic and aposymbiotic samples were analyzed using Seurat (v.5.0.2) (46). Datasets were cleaned (genes expressed in fewer than three cells were discarded, as were cells expressing fewer than 200 genes or greater than 3,000 genes), normalized, and scaled, and a PCA reduction was performed. No duplicated barcodes were detected. Mitochondrial reads could not be removed, as the reference genomes and transcriptome do not have annotated mitochondrial genomes. Datasets were integrated using the Canonical Correlation Analysis (CCA) Integration method on the PCA reduction on the top 2,000 most variable genes (nearest neighbors parameters). Cell clusters were identified using 30 dimensions (resolution: 0.5) and visualized using Uniform Manifold Approximation and Projection (UMAP). Marker genes for each cluster were identified using FindAllMarkers (Wilcoxin Rank Sum Test; log_2_foldchange threshold: 0.5) and cell types were informed by gene annotations and marker gene comparisons from other cnidarian single-cell datasets (e.g., *S. pistallata* and *Xenia* sp.) (**Supplementary Dataset 2**) (47,48). Expression patterns of top marker genes were assigned to cell clusters using violin plots, bubble plots, and visualization of gene expression in individual cells within the UMAP. 28 cell clusters across 7 cell types were identified in the UMAP. Broad transcriptomic differences between cell clusters were assessed by identifying the top 10 genes enriched in each cluster compared to all the other clusters.

To identify cells containing algal symbionts, we identified cells in which over 50% of the total reads corresponded to *B. psygmophilum* genes, and then visualized these cells on the UMAP. These cells were then removed, as the low percentage of host reads in these cells confounded downstream analyses.

To determine how symbiosis alters enrichment of gene pathways at the whole-organism level, and how this differential enrichment is reflected in specific cell types, we performed Mann– Whitney U Gene Ontology (GO) term enrichment analysis on all genes from the full single-cell dataset (49). Significantly enriched GO terms within the ‘Biological Process’ (BP) GO division were defined with a false-discovery rate (FDR) of <0.1. Significantly over- and under-represented GO terms relating explicitly to immunity, antigen signaling, or the NF-κB pathway were identified (**Table S1**). Cells in which over 7% of the total reads corresponded to genes annotated with the GO terms of interest were visualized on the UMAP of all cells from symbiotic and aposymbiotic *O. arbuscula* samples.

To compare expression profiles of specific cell states between symbiotic and aposymbiotic samples, each cell cluster was independently reclustered, and the effect of symbiotic state on gene expression in each cell state was analyzed. First, the cell cluster of interest (*e.g.,* Immune Cell, Gastrodermis 1) was subsetted and new variable features were identified. Data were rescaled, and dimensionality reduction was re-run (first 30 dimensions). Subclusters were identified with resolutions between 0.25-0.5 (**Table S2**). The distribution of cells across subclusters between symbiotic states was visualized using UMAPs. Differentially expressed genes between symbiotic states in all cell clusters were identified using DESeq2 within the FindMarkers function. Additionally, marker genes for each subcluster were identified. To highlight differences in the gene expression of the Immune Cell states, the top 20 genes enriched in each Immune Cell subcluster were identified and plotted (unannotated genes were removed).

We used Monocle 3 (v.1.3.7) (50) to determine cell trajectories and outcomes for all gastrodermal cells from symbiotic and aposymbiotic samples. We plotted the UMAPs of the symbiotic and aposymbiotic subsets of Gastrodermis clusters 1-4 and overlaid trajectory graphs, noting nodes and outcomes.

To compare expression of genes involved in sugar transport and nutrient cycling in gastrodermal cells between symbiotic aposymbiotic states, the expression of these genes in the reclustered Gastrodermis 1 and Gastrodermis 2 cells was analyzed. First, normalized expression of genes involved in nitrogen cycling/symbiont density control (Glutamate Dehydrogenase DHE4 and Glutamate Synthase [NADH] GLSN) and sugar transport (Apolipophorin APOB) (14) were compared across all cell clusters using violin plots (function VlnPlt in the Seurat package (46)). FeaturePlot (46) was then used to visualize co-expression of these genes across the subclusters of Gastrodermis 1 and Gastrodermis 2 cells from the symbiotic sample (scaling per-cell mean expression to a maximum value of 10). To determine whether the coexpression of genes observed in the symbiotic gastrodermal cells is seen in other cell types, we also visualized coexpression of DHE4, APOB, and GLSN across Epidermis 1 and Neuron 1 from symbiotic tissue (scaling per-cell mean expression to a maximum value of 10).

To identify genes driving separation between symbiotic states within each cell cluster, the distribution of cells from a particular cluster was visualized along the first 10 PCs of the PCA assay used to define the UMAP for that cluster. An analysis of variance determined whether the cells were separated by symbiotic state (symbiotic vs. aposymbiotic) along each PC. In Gastrodermis 1 and Gastrodermis 2 clusters, the top 20 genes driving the distribution of cells along each significant PC (10 genes driving each PC in the positive direction and 10 genes driving each PC in the negative direction) were selected, and unannotated genes were removed. In Gastrodermis 1, PCs 1, 3-8, and 10 were selected for a total of 96 genes. In Gastrodermis 2, PCs 1, 2, 5, 7, and 8 were selected for a total of 59 genes. Log_2_foldchange values from DESeq2 with accompanying *p*-adjusted values (<0.05) for each gene were plotted in a bar graph. Genes annotated with immune-function related GO terms were noted, as were genes annotated with the Clusters of Orthologous Genes (COG) annotation ‘posttranslational modification, protein turnover, and chaperones.’

Finally, to compare *O. arbuscula* cells potentially hosting algal symbionts with algal-hosting cells from an obligate coral species (*Xenia* sp. (48)), we identified orthologous genes between *A. poculata* (the mapping reference in this study) and *Xenia* using the Broccoli algorithm using 13 cnidarian proteomes with default parameters (51). We re-ran Seurat normalization, PCA, and UMAP clustering on the pre-integrated *Xenia* dataset from non-regeneration samples (48) to identify 16 clusters (resolution of 0.18 and 30 dimension), including the previously characterized algal-hosting cell cluster and the other two gastrodermal cell clusters. We identified differentially expressed genes between the algal-hosting cells and the gastrodermal cells (*p*-adjusted <0.05 from the FindMarkers function, using both gastrodermal cell clusters as the comparison to the algal-hosting cell cluster), and subsetted these genes for those that had orthologs to *A. poculata*. Additionally, we identified differentially expressed genes between algal-hosting and non-algal hosting *O. arbuscula* cells in two ways: 1) from symbiotic *O. arbuscula* cells from Gastrodermis 1/Gastrodermis 2 and aposymbiotic *O. arbuscula* cells from Gastrodermis 1/Gastrodermis 2 (*p*-adjusted <0.05 from the FindMarkers function), and 2) from symbiotic *O. arbuscula* cells from Gastrodermis 1/Gastrodermis 2 and from symbiotic *O. arbuscula* cells from Gastrodermis 3/Gastrodermis 4. We subsetted these genes to include only *Xenia* orthologs. We compared which genes among these three sets of orthologs (*Xenia*-sym-to-gastro, *Oculina-*sym-to-apo, and *Oculina*-sym-to-gastro respectively) were differentially expressed using a Venn diagram (52), creating six groups of differentially expressed orthologs (DEOs). We performed Fisher Exact Tests (DEO presence/absence) (49) to identify enrichment differences across these groups of DEOs in the BP GO division.

## RESULTS

For the following experiments, genetically controlled symbiotic and aposymbiotic fragments of *O. arbuscula* were used. Symbiotic fragments had symbiont densities that were 400-fold greater (13,678 +/− 6,204 cells/cm^2^) than those of aposymbiotic fragments (30 +/− 43 cells/cm^2^) (p<0.001) (**Fig. 1a**). These fragments were then used to generate light-scattering profiles, 16S metabarcoding libraries, tissue proteomics, and single-cell profiles.

**Figure 1:**
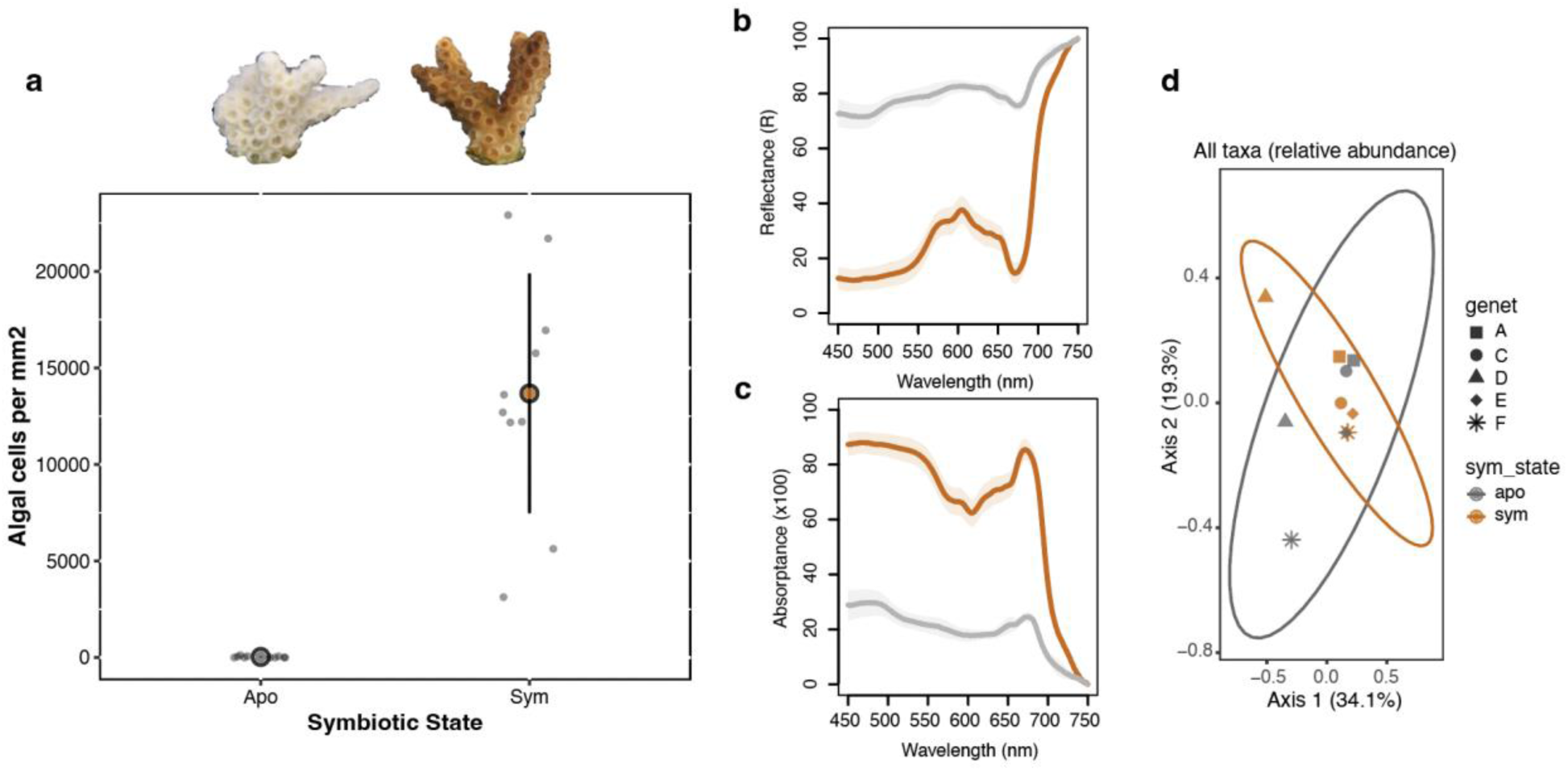
Symbiosis facilitates higher light absorptance but no changes to bacterial communities in *O. arbuscula*. **a** Algal cell counts are 400x higher in symbiotic (brown, right) *O. arbuscula* fragments than in aposymbiotic (white/gray, left) fragments (p<0.001). **b** Average reflectance (R) in the PAR region (400–700 nm) was 22% +/− 4% in symbiotic *O. arbuscula* (brown) and 77% +/− 3% in aposymbiotic *O. arbuscula* (gray) (p < 0.01) **c** Average absorptance (A) (A = 1-R) demonstrating relative amount of solar energy with potential for use in photosynthesis. Symbiotic *O. arbuscula* (brown) has an A_675_ of 85% (+/− 4%), suggesting a functional symbiosis, and aposymbiotic (gray) has an A_675_ of 24% (+/− 2%) (p<0.001). In **b** and **c**, shaded colored areas represent standard deviation. **d** PCoA of bacterial communities from all ASVs in 16S data from symbiotic (brown) and aposymbiotic (gray) *O. arbuscula* genotypes. No differences in bacterial communities were observed between symbiotic states.

### Light absorption is higher in symbiotic *O. arbuscula*

Differences in symbiont cell densities induced significant differences in *in vivo* coral light absorption. Reflectance spectra measurements and absorption properties of intact corals showed statistically significant pigment content differences between symbiotic and aposymbiotic *O. arbuscula*. Average reflectance in the PAR region (400–700 nm) of symbiotic fragments was 22% +/− 4%, whereas aposymbiotic coral reflectance was 77% +/− 3% (p < 0.01) (**Fig. 1b**). In symbiotic fragments, the absorptance value at the peak for Chlorophyll a (A_675_) was 85% +/− 4%, compared to 24% +/− 2% in aposymbiotic samples (p<0.0001) (**Fig. 1c**). Absorptance describes the relative amount of solar energy/incident light that can potentially be used in photosynthesis for organic carbon fixation.

### Bacterial communities do not differ across symbiotic states

We profiled the bacterial communities of aposymbiotic and symbiotic fragments from 5 genets (N=10). A total of 208,330 sequences were acquired with a mean depth of coverage of 20,833 +/− 16050 per sample (**Table S3**). Sample rarefication yielded a total of 8,298 reads per sample with a total of 661 ASVs identified across samples. No differences in bacterial communities were observed between symbiotic and aposymbiotic fragments (**Fig. 1d**), and both states were dominated by Proteobacteria and Bacteroidota (**Fig. S1**). Additionally, no differences in alpha diversity were observed between symbiotic states regardless of the metric tested, nor were differences in beta diversity by symbiotic state detected (**Fig. S2a-d**). DESeq2 tested for differentially abundant ASVs across symbiotic states, while modeling genetic background, but no ASVs were identified.

### The proteome is affected by symbiotic state in *O. arbuscula*

To compare proteomes of symbiotic and aposymbiotic *O. arbuscula*, we performed mass spectrometry on whole-cell lysates from coral fragments. We identified a total of 2,343 proteins in *O. arbuscula*, 2,191 of which were retained following filtering. A PCA showed that samples clustered by symbiotic state (*p_state_*<0.1) and genet (*p_genet_*<0.05) (**Fig. 2a**). A total of 187 proteins were differentially enriched across symbiotic states (*p-*adjusted<0.01: 105 enriched in symbiotic, 82 enriched in aposymbiotic *O. arbuscula*; **Figs. 2b-c**).

**Figure 2:**
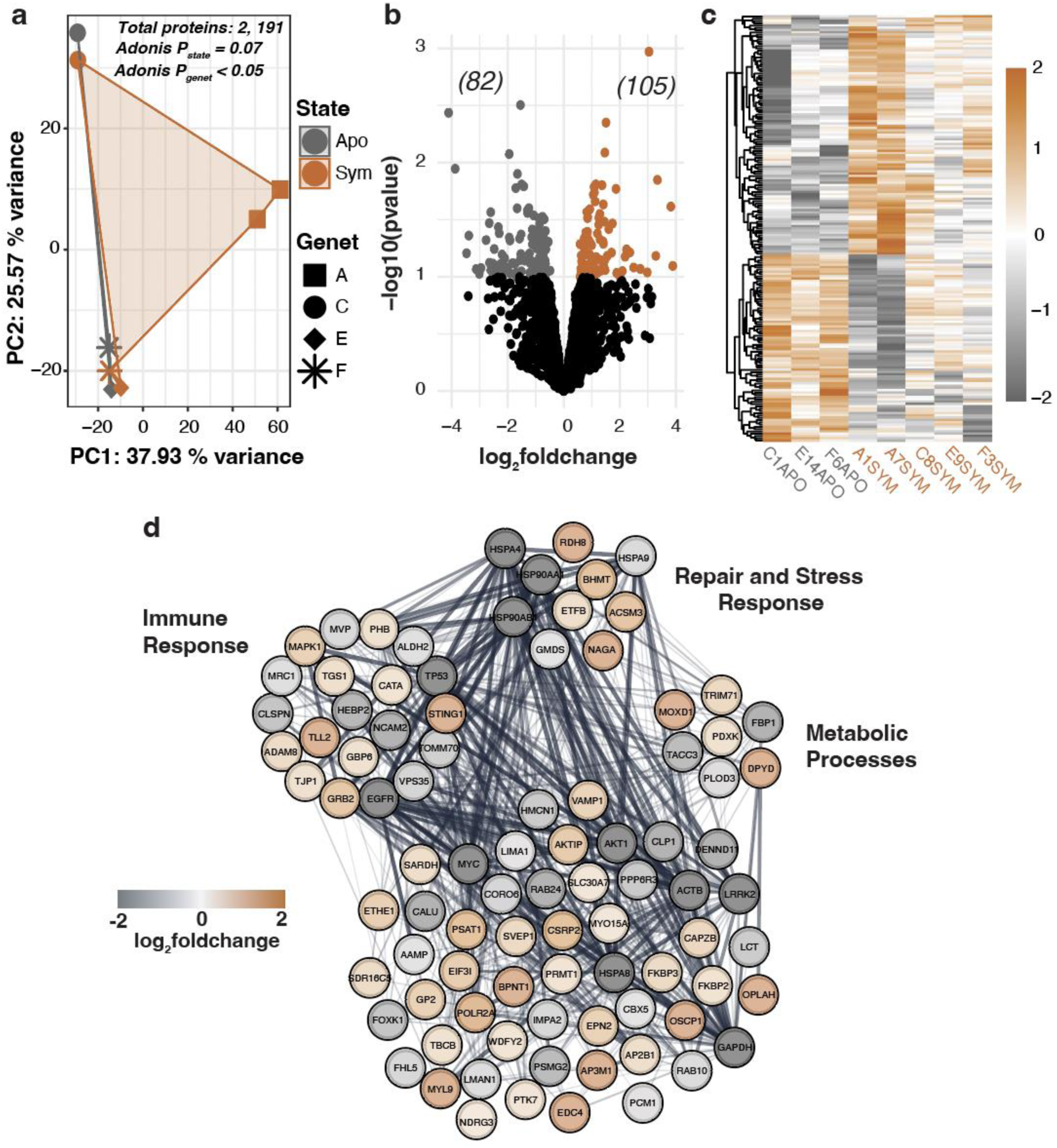
Proteomic profiles of symbiotic and aposymbiotic *O. arbuscula* reveal differential enrichment of proteins by symbiotic state. **a** Principal component analysis of proteomic profiles across four genets, with symbiotic (sym) and aposymbiotic (apo) states distinguished by color, and genets by shape. **b** Volcano plot of differentially enriched proteins (DEPs) identified through pairwise comparison (*p*-value < 0.1). Upregulated proteins (105) are represented by brown dots, downregulated proteins (82) by gray dots, while black dots indicate non-DEPs in symbiotic relative to aposymbiotic corals (total N=2,191). **c** Heatmap of all DEPs across symbiotic states of *O. arbuscula*. The color scale represents the log_2_foldchange of each protein (row) for each coral sample (column) relative to the protein’s mean across all samples. **d** Interaction network of proteins associated with Immune Response, Repair and Stress Response, and Metabolic Processes, derived from human protein–protein interactions in the STRING database. Node colors indicate upregulation (brown) or downregulation (gray) in symbiotic coral samples.

Differentially enriched proteins (DEPs) were assigned functional categories based on a homology search of each DEP sequence in the UniProtKB database (evalue < 1 × 10^-5^). DEPs involved in Immune Response, Repair and Stress Response, and Metabolic Processes were identified (**Fig. 2d**)

### Single-cell RNA-seq identifies cell states shared between symbiotic and aposymbiotic *O. arbuscula*

To profile gene expression in different cell types and states from symbiotic and aposymbiotic *O. arbuscula*, we used the 10X single-cell RNA-sequencing (scRNA-seq) platform. We captured a total of 6,992 cells (3,158 cells from the symbiotic sample; 3,834 cells from the aposymbiotic sample), with an average of 157,534.5 reads per cell, 1,121 median UMI counts per cell, and 651.5 median genes per cell. The average number of reads was 540,436,426; the average percentage of valid barcodes was 94.85%; the average percentage of valid UMIs was 99.90%; the average sequencing saturation was 89.75%. Further quality control data for both samples are summarized in **Supplementary Dataset 1**.

Clustering of the integrated 10X gene expression datasets from symbiotic and aposymbiotic samples resulted in the identification of 28 transcriptomically distinct cell clusters across seven broad cell types (**Fig. 3a**). Cells from symbiotic and aposymbiotic samples were represented in all clusters (**Fig. 3a**). Additionally, each cluster defined in the UMAP displayed distinct transcriptomic programming as exemplified by expression patterns of the ten highest enriched genes in each cluster (**Fig. 3b**).

**Figure 3:**
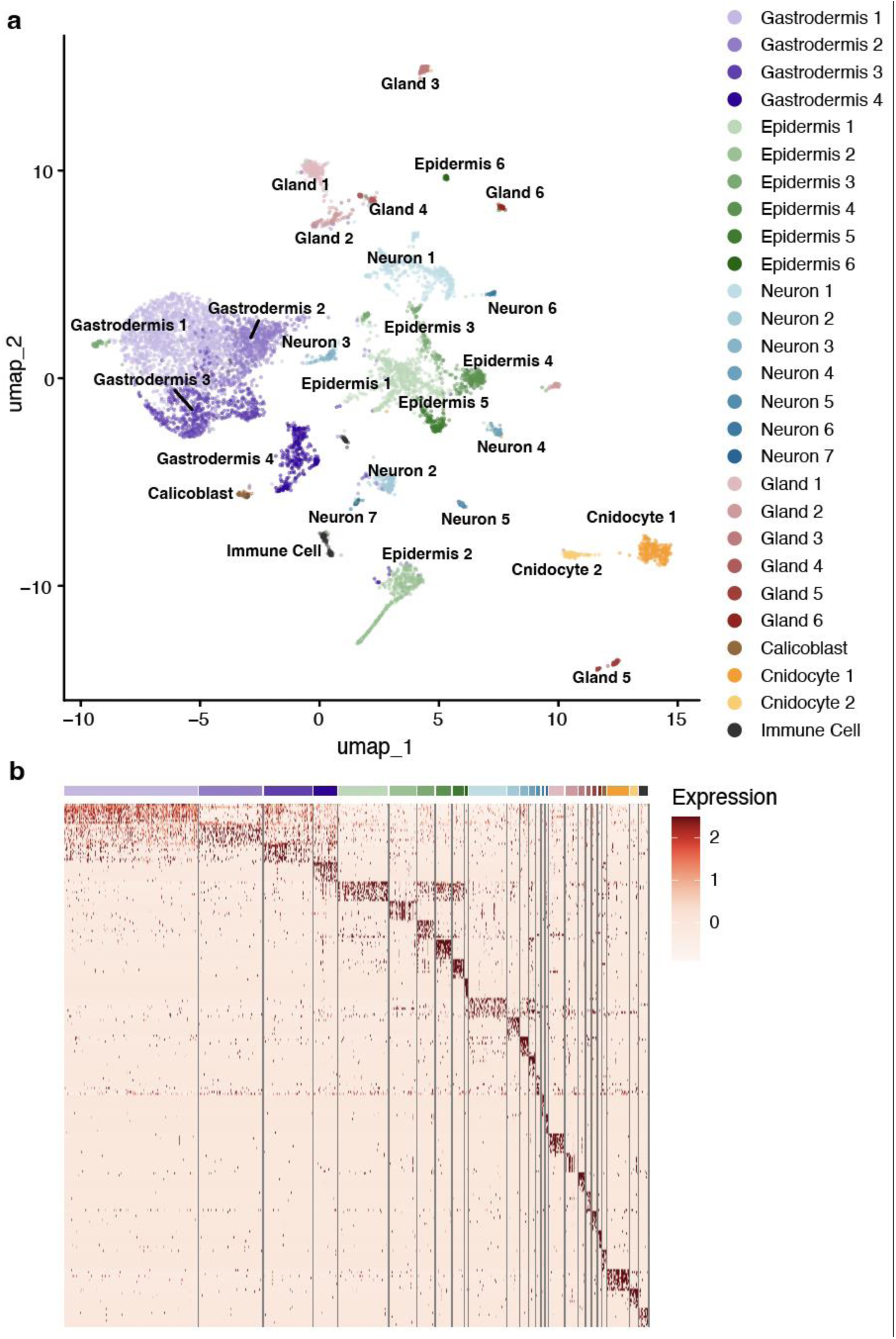
scRNA-seq reveals 28 transcriptomically distinct cell clusters across 7 cell types in symbiotic and aposymbiotic *O. arbuscula*. **a** A UMAP Projection of the transcriptomes of 6,992 cells from symbiotic (sym) and aposymbiotic (apo) *O. arbuscula* reveals 28 clusters across 7 broad cell types. Each cluster is color coded. **b** Each cluster is transcriptomically distinct, with expression similarities present between clusters of the same cell type, as represented by the 10 genes most highly expressed in each cluster. Expression values are average log_2_foldchanges computed from normalized gene expression data for each cell. Cell clusters are colored as in (**a**).

### Identification of symbiont-hosting cells

We identified 34 cells in which over 50% of the total reads corresponded to *B. psygmophilum* genes. 33 of these 34 cells belonged to the symbiotic Gastrodermis 2 cluster (**Fig. S3**), indicating that this cell cluster contained the algal-hosting cells from symbiotic *O. arbuscula*.

### Symbiotic gastrodermal cell populations fated for symbiosis function in nutrient exchange

To investigate the cell fates of the gastrodermal cells, we performed trajectory analysis on the four gastrodermal cell clusters in the symbiotic and aposymbiotic *O. arbuscula* samples separately. In the symbiotic sample, we identified two fates, one terminating in Gastrodermis 2 where symbiont-hosting cells were identified, and one terminating in Gastrodermis 3 (**Fig. 4a**). One fate was identified in the aposymbiotic gastrodermal cells, terminating in Gastrodermis 4 (**Fig. S4**). To probe gene expression differences between gastrodermal cells from symbiotic compared to aposymbiotic *O. arbuscula*, we subsetted and reclustered cells from each of the four gastrodermal states. Four subclusters were revealed within Gastrodermis 1 (**Fig. 4b**), three subclusters were revealed within Gastrodermis 2 (**Fig. 4c**), two were revealed in Gastrodermis 3 (**Fig. 4d**), and one was revealed within Gastrodermis 4 (**Fig. S5a**), suggesting additional cell states. Gastrodermis 1 Subcluster 3 and Gastrodermis 2 Subcluster 3 were composed almost entirely of cells from the symbiotic sample (**Fig. 4b** and **Fig. 4c** respectively). Independent reclustering of all other cell types was also performed, and the resulting UMAPs displayed varying, though lesser, degrees of structure by symbiotic state (**Fig. S5b-w**).

**Figure 4:**
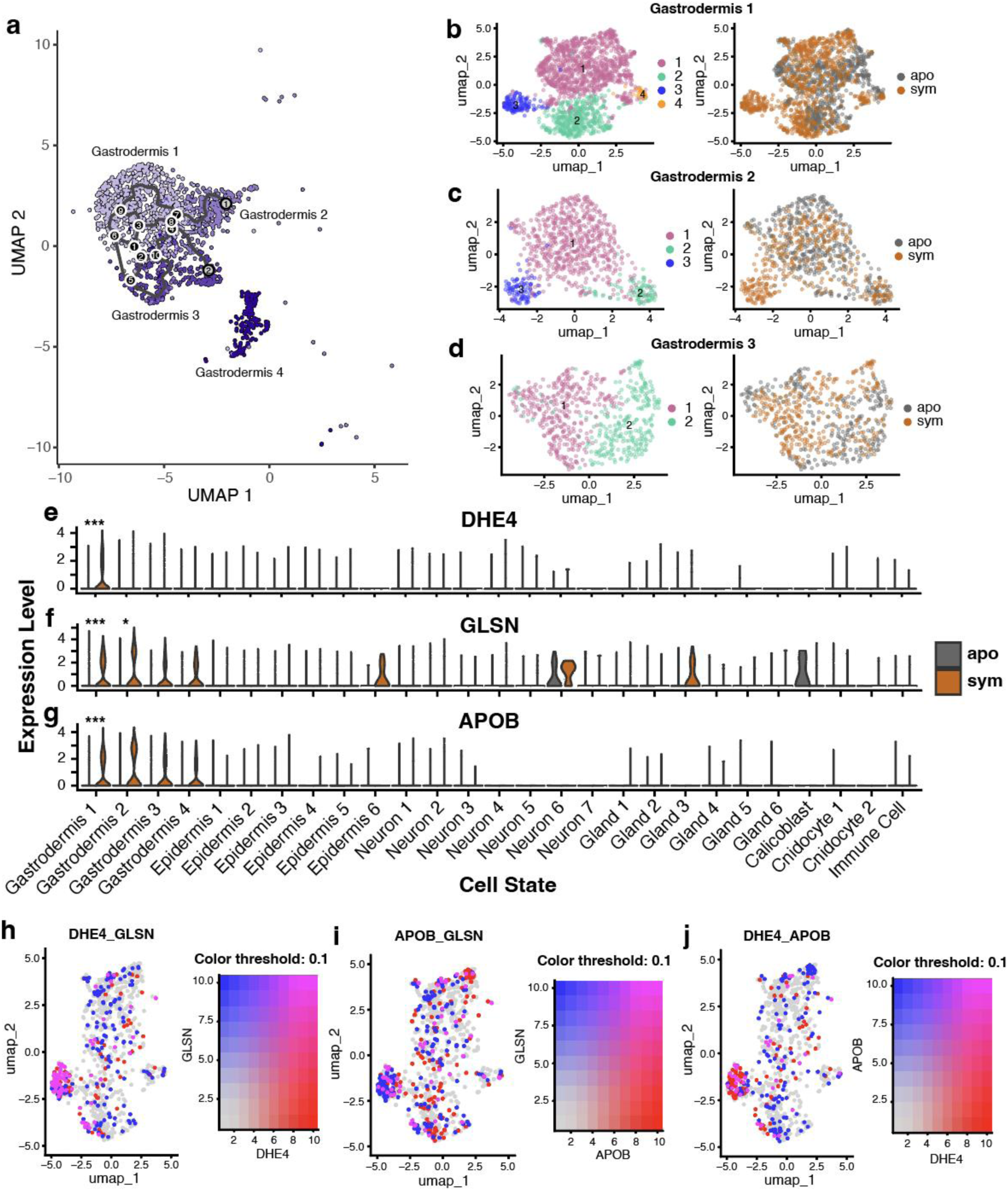
Gastrodermis cells along the algal-hosting trajectory are involved in sugar transport and nitrogen cycling. **a** UMAP projection of Gastrodermis 1, 2, 3, and 4 cells from symbiotic *O. arbuscula* overlayed with a graph of cell trajectories. Outcomes (fates) are denoted by gray circles with black numbers. Branch nodes are denoted by black circles with white numbers. **b** UMAP projections of the 1,682 independently reclustered Gastrodermis 1 cells revealed 4 subclusters. Each subcluster is color-coded in the left panel. Gastrodermis 1 cells separate according to symbiotic state across the UMAP projection (right panel). **c** UMAP projections of the 802 independently reclustered Gastrodermis 2 cells revealed 3 subclusters, color coded in the left panel. Gastrodermis 2 cells separate according to symbiotic state (right panel). **d** UMAP projections of the 611 independently reclustered Gastrodermis 3 cells revealed 2 subclusters, color coded in the left panel. Limited separation of Gastrodermis 3 cells by symbiotic state (right panel) was observed. Violin plots of Glutamate Dehydrogenase (DHE4, **e**), Glutamate Synthase [NADH] (GLSN, **f**), and Apolipophorin (APOB, **g**) across all cell clusters in both symbiotic states. DHE4 was identified as a marker gene for Gastrodermis 1; GLSN and APOB were identified as marker genes for both Gastrodermis 1 and 2. Asterisks denote genes differentially expressed in symbiotic versus aposymbiotic clusters (* = *p*-adjusted value <0.05; *** = *p*-adjusted value < 0.001). Expression levels of genes in **e-f** are normalized per cell, and significance is calculated from DESeq2. Coexpression plots of DHE4 and GLSN (**h**), APOB and GLSN (**i**), and DHE4 and APOB (**j**) across Gastrodermis 1 cells from symbiotic *O. arbuscula* revealed high coexpression of these sugar transport and nitrogen cycling genes within Subcluster 3, but not in the other subclusters. The coloration of each cell represents the per-cell mean expression value of each gene scaled to a maximum value of 10.

To help identify the localization of the gastrodermal cells participating in active symbiotic nutrient exchange within our dataset, we selected sugar transport and nitrogen cycling genes, which have been previously shown to have higher expression in symbiotic compared to aposymbiotic *O. arbuscula* (14). DESeq2 identified differences in gene expression in gastrodermal cells between symbiotic and aposymbiotic samples, which showed that Gastrodermis 1 cells from symbiotic *O. arbuscula* have higher expression of genes related to nitrogen cycling (Glutamate Dehydrogenase DHE4 and Glutamate Synthase [NADH] GLSN) and sugar transport (Apolipophorin APOB) (*p*-adjusted <0.05) (**Fig. 4e-g**). Furthermore, DHE4, GLSN, and APOB were identified as marker genes for Subcluster 3 (indicating that these genes were more highly expressed inside than they were outside Subcluster 3 (*p*-adjusted <0.05). Additionally, there was strong co-expression of these symbiosis-marker genes in Subcluster 3, suggesting that this cell state is actively transporting fixed carbon sugars while mediating the growth and division of algal symbionts (**Fig. 4h-j**). There was less co-expression of these genes in Subclusters 1, 2, and 4 of Gastrodermis 1, suggesting that other gastrodermal cell states perform nitrogen and carbon sugar processing independently or through distinct pathways. Similar, though weaker, patterns of coexpression were observed in Gastrodermis 2 (**Fig. S6a-c**). Additionally, there was no or limited coexpression of DHE4, APOB, and GLSN observed in symbiotic Epidermis 1 and Neuron 1 cells (**Fig. S6d-i**). The presence of symbiotic-only subclusters within Gastrodermis 1 and 2, coupled with the co-expression of highly expressed markers for symbiosis within one Gastrodermis 1 subcluster suggest that certain gastrodermal cells states in our dataset are actively hosting symbionts.

### No differences in immune cell gene expression between symbiotic state

To determine whether downregulation of genes associated with the immunity observed in whole-organism studies of symbiotic compared to aposymbiotic coral (this study and (14)) was observed in the Immune Cell gene expression, we identified one Immune Cell cluster with high expression of cell markers for immunity. These immune marker genes included Interferon Regulatory Factors (IRFs), Activating Transcription Factor 3 (ATF3), and multiple chaperone proteins in cells from both symbiotic and aposymbiotic tissue (**Fig. 5a, b**). It is worth noting that we did not identify cell type- or symbiotic state-specific expression of other commonly studied cnidarian immune genes, including NF-κB, cGAS, and STING (20,53). After subsetting and independently reclustering these Immune Cells, we identified three cell states with distinct transcriptomic profiles (**Fig. 5c**). However, cells did not cluster by symbiotic state after reclustering, and no differential expression between Immune Cells from symbiotic and aposymbiotic samples was observed (*p*-adjusted <0.05; **Fig. 5d**).

**Figure 5:**
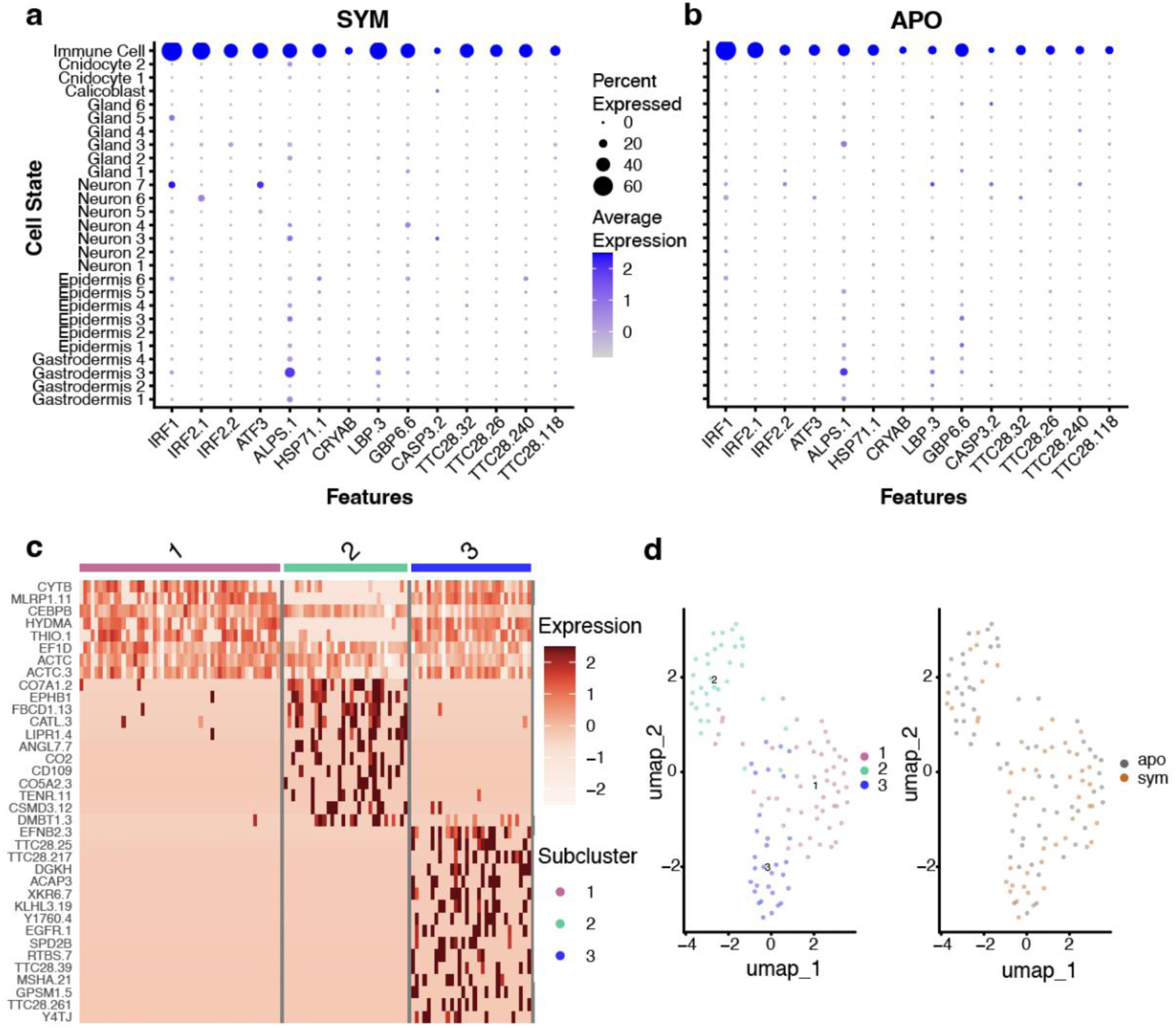
Negligible gene expression differences between Immune Cells of symbiotic and aposymbiotic samples. Immune Cells from symbiotic (**a**) and aposymbiotic (**b**) samples have high expression of previously identified immune cell marker genes. Dot size represents the percent of cells in each cell state that express each gene, and dot color represents the average (normalized) expression of each gene within each cell state. **c** Three Immune Cell sublcusters have distinct transcriptomic profiles, as represented by the annotated genes within the top 20 most highly enriched genes in each subcluster. Expression values are average log_2_foldchanges computed from normalized gene expression data of each cell. **d** UMAP projections of the 115 independently reclustered Immune Cells revealed 3 subclusters (left panel) that do not separate according to symbiotic state (right panel).

### Immune response genes are differentially regulated in symbiotic Gastrodermis 1 and 2 cells

In agreement with previous bulk studies (14,20,54), our dataset demonstrated that, at the whole-organismal scale, immune system gene pathways identified through GO term analysis are differentially regulated under symbiosis. Interestingly, genes belonging to these differentially enriched immunity GO terms are most highly expressed in gastrodermal cells. In addition, expression of these genes is higher in gastrodermal cells from aposymbiotic (**Fig. 6a**) compared to symbiotic *O. arbuscula* (**Fig. 6b**), demonstrating that symbiont-specific immune gene regulation occurs in gastrodermal cells rather than immune cells. Furthermore, because Gastrodermis 1 and 2 cells clustered by symbiotic state, we identified the factors (genes) driving this pattern in the Gastrodermis 1 and 2 cell UMAPs. We analyzed the distribution of the cells along each of the first 10 Principal Components (PCs) in the PCA dimensionality reduction used to generate the UMAPs. Gastrodermis 1 cells exhibited significant separation by symbiotic state along 8 of the first 10 PCs (*p*-value < 0.05), and Gastrodermis 2 cells showed significant separation along 5 of the first 10 PCs. In Gastrodermis 1, of the 96 annotated genes driving the distribution of the cells along the 8 significant PCs, 33 were differentially expressed (*p*-adjusted <0.05), and 32 of the 96 genes were annotated with GO terms involved in immunity (**Fig. 6b**). Additionally, 9 genes were annotated with the COG term ‘posttranslational modification, protein turnover, and chaperones’ (**Fig. 6b**). In Gastrodermis 2, 4 of the 59 genes were differentially expressed (*p*-adjusted <0.05), 13 were annotated with immunity GO terms, and 5 were annotated with the COG term ‘posttranslational modification, protein turnover, and chaperones’ (**Fig. 6c**).

**Figure 6:**
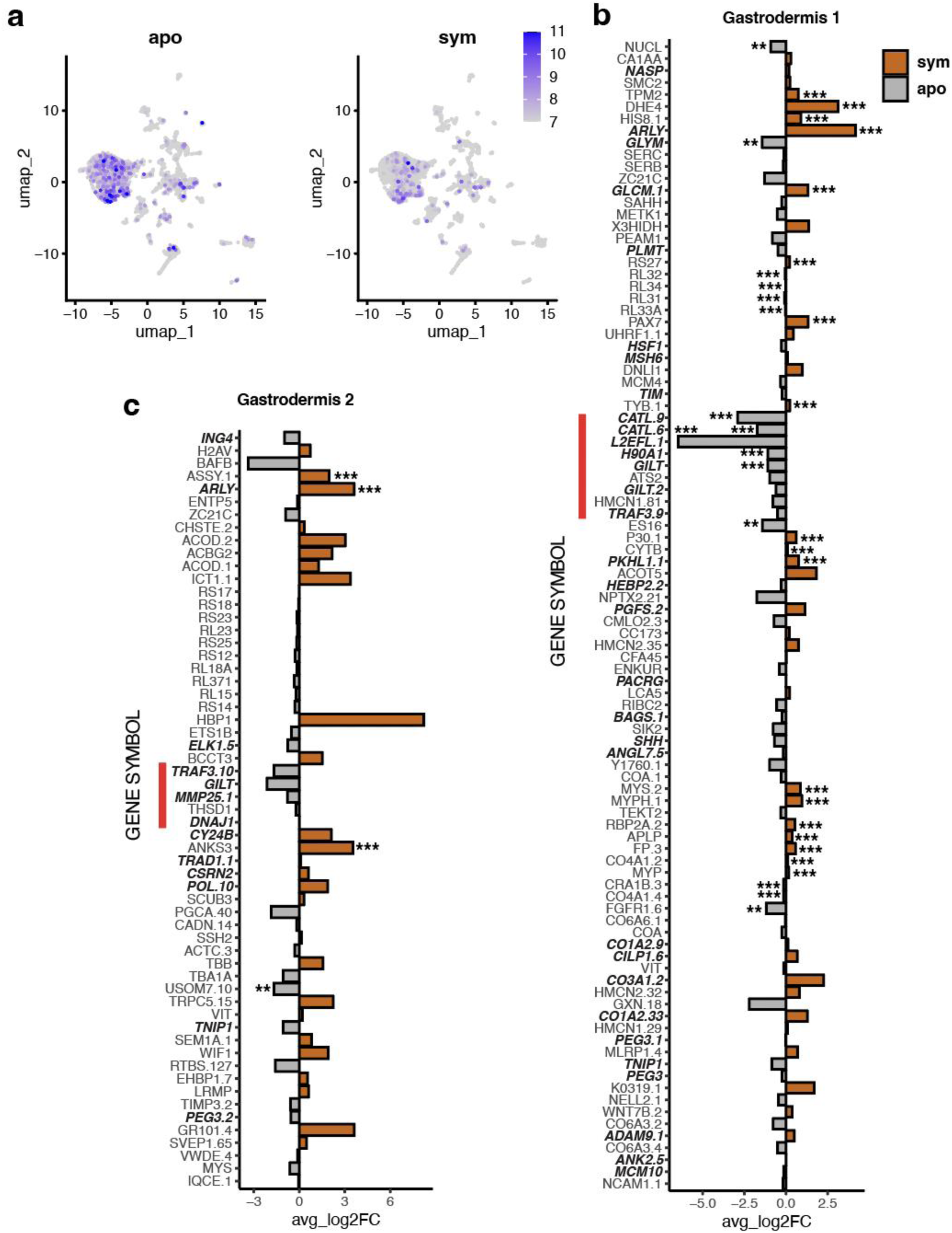
Gastrodermis 1 and 2 cells from symbiotic samples downregulate immune pathway genes compared to aposymbiotic samples. **a** Spatial representation showing expression of genes annotated with differentially enriched immunity GO terms across the whole dataset. Coloration represents the percentage of all transcripts in each cell corresponding to the genes of interest (555 genes of interest out of 24,716 total genes). **b** 8 PCs, driven by 96 annotated genes, contributed to the separation of symbiotic state in the reclustered UMAP of Gastrodermis 1 (see Fig. 4b). 33 of the 96 were differentially expressed. 32 genes are annotated with GO terms relating to immunity (bold, italicized). **c** 5 PCs, driven by 59 annotated genes, contributed to the separation of symbiotic state in the reclustered UMAP of Gastrodermis 2 (see Fig 4c). 4 of the 59 genes were differentially expressed. 13 genes are annotated with GO terms relating to immunity (bold, italicized). In both **b** and **c**, the red bar indicates genes annotated with the COG term ‘posttranslational modification, protein turnover, and chaperones.’ Significance is calculated from DESeq2 (*** = *p*-adjusted value < 0.001, *** = *p*-adjusted value < 0.01, * = *p*-adjusted value < 0.05).

### In two coral species, gastrodermal cells capable of symbiosis share functions relating to metabolism

To compare conserved mechanisms by which specific types of gastrodermal cells host algae across coral species, we analyzed the overlap of differentially expressed orthologs (DEOs) between algal-hosting cells (Gastrodermis 1 and 2 cells along the same cell fate trajectory) and non-algal-hosting gastrodermal cells from our facultatively symbiotic coral to algal-hosting cells and other gastrodermal cells from an obligate coral species (*Xenia* sp. (48)). We identified 200 DEOs between *Xenia* algal-hosting and non-algal hosting gastrodermal cells (*Xenia*-sym-to-gastro), 233 DEOs between *O. arbuscula* Gastrodermis 1 and 2 cells from symbiotic and aposymbiotic tissue (*Oculina-*sym-to-apo), and 184 DEOs between *O. arbuscula* Gastrodermis 1 and 2 cells from symbiotic tissue and Gastrodermis 3 and 4 cells from symbiotic tissue (*Oculina*-sym-to-gastro). We performed this analysis in two ways to account for the facultatively symbiotic nature of *O. arbuscula*. First, we analyzed overlapping and unique DEOs between *Xenia*-sym-to-gastro and *O. arbuscula Oculina-*sym-to-apo: 71 DEOs were shared, 129 were unique to *Xenia*-sym-to-gastro, and 162 were unique to *Oculina-*sym-to-apo (**Fig. S7a**). Second, we analyzed overlapping and unique DEOs between *Xenia*-sym-to-gastro and *O. arbuscula Oculina-*sym-to-gastro: 51 DEOs were shared, 149 were unique to *Xenia*-sym-to-gastro, and 133 were unique to *Oculina-*sym-to-gastro (**Fig. S7b**). Using a Fisher Exact Test of BP GO terms, we probed functional enrichment of processes across each DEO group from the Venn diagrams. Across shared and *Oculina*-unique DEO groups, the most common GO terms were involved in metabolism, protein localization, and ribosomal function. Interestingly, the functions relating to cellular stress (e.g., “Cellular Response to DNA Damage Stimulus”) and cell cycle regulation (e.g., “Regulation of Cell Cycle”) were enriched in *Oculina*-sym-to-gastro-unique DEO set; furthermore, the “Immune Response” GO term was enriched in both *Xenia*-sym-to-gastro-unique DEO sets (**Supplementary Dataset 3**).

## DISCUSSION

We characterized how host immunity is modulated to facilitate endosymbiosis by quantifying phenotypic metrics of symbiosis and profiling bacterial communities, whole-organism proteomics, and single-cell RNA sequencing (scRNA-seq) in symbiotic (with algal symbionts) and aposymbiotic (without algal symbionts) samples of the facultatively symbiotic stony coral *Oculina arbuscula.* This work reveals cell type-specific compartmentalization of the immune system that allows symbiotic hosts to simultaneously downregulate key immune pathways in algal-containing cells while maintaining constitutive organismal immunity across other cell types. Our results are consistent with emerging data indicating that the cnidarian-algal symbiosis is a balancing act between nutritional status and immunity (14,20,55). Hosts must balance beneficial nutrient acquisition from photosynthetic algae with the downregulation of the immune system to allow for intracellular symbiont establishment and maintenance.

Whole-organism transcriptomic and proteomic research has yielded many insights into how symbiosis influences cnidarian host functions. For example, transcriptomic (14) and proteomic ((55) and this study) experiments have demonstrated how symbiotic state affects gene expression and protein abundance. Our whole-organism proteomic data across different symbiotic states under baseline conditions support results from previous experiments (14,20,54) demonstrating that differential regulation of the host immune system is required to maintain symbiosis. Specifically, our whole-organism proteomic analysis and bulk analysis of our single-cell data confirmed the pattern of differential regulation of immune system pathways under symbiosis in a facultatively symbiotic cnidarian, which has been previously observed in *O. arbuscula* (14), *E. pallida* (20) and *A. poculata* (27). However, signals from whole-organism studies may be confounded by differences in the proportions of cell types sampled and opposing expression signals within these cell types (56). Furthermore, while observed patterns between scRNAseq and proteomic profiling are consistent between aposymbiotic and symbiotic fragments, we cannot determine whether the signature in the bulk proteomic data is driven by gastrodermal algal-hosting cells. Future work coupling scRNAseq with single cell proteomic profiling (57) or immunohistochemistry would reveal more information about the localization of differentially enriched proteins across cell types.

scRNA-seq has been gaining traction in research on the coral immune system. In one study, Levy et al. characterized two immune cell states with divergent expression patterns in obligate symbiotic stony coral *Stylaphora pistillata* adults (47). In additional studies, Hu et al. described how LePin, a lectin involved in the phagocytic pathways of the cnidarian immune system, is required for symbiont establishment in algal-hosting cells of the soft coral *Xenia* sp. (48,58). Cell state-specific regulation of the cnidarian immune system has also been proposed to occur in the sea anemone *E. pallida* via whole-larvae fluorescence imaging and transcriptomics (59). Notably, MYD88, a protein that transduces extracellular signals into the host to induce the NF-κB pathway (60,61) is repressed in *E. pallida* algal-hosting cells; this local immune modulation only occurs in symbiosis with compatible algae, thereby allowing host cells to sort and expel other non-symbiotic microbes (59). Our scRNAseq approach further highlights how the *O. arbuscula* immune system is compartmentalized and reveals how cell state-specific immune modulation may be key to the maintenance of a functionally symbiotic host. Here, we observe a downregulation of immune pathway genes in populations of symbiotic gastrodermal cells along the cell fate trajectory that terminates in algal-hosting cells, indicating that immune suppression is key to symbiosis maintenance in these cell types. Although we were unable to confidently identify LePin in our data, we noted the clear involvement of immune gene regulation in driving transcriptomic differences in gastrodermal cells from symbiotic and aposymbiotic tissue. Notably, these genes included Heat-Shock Proteins (HSPs) and Cathepsins, which can be involved in the NF-κB pathway and protein chaperoning and are hypothesized to function in symbiosis regulation (62–65). Induction of HSPs has been observed following thermal stress in symbiotic cnidarians, implicating the unfolded protein response in ensuring the stability of symbiosis (63,66). Interestingly, HSPs and Cathepsins have been previously shown to be markers for immune cells in the stony coral *S. pistillata* (47), but it remains to be determined whether this immune signature in gastrodermal cells is conserved in cnidarians engaged in obligate symbioses. It is important to note that of the two gastrodermal cell types that we identified as displaying symbiosis-induced immune gene suppression, Gastrodermis 1 exhibited stronger signals than Gastrodermis 2. This observation is most likely due to the fact that Gastrodermis 1 contains more than twice the number of cells as Gastrodermis 2 (1,682 cells versus 802 cells) and therefore had more statistical power. Evidence for localized regulation of immune system pathways in algal-hosting gastrodermal cells has now been shown in a soft coral (58), a sea anemone (59), and a stony coral (present study). This compartmentalized immune system regulation allows for the maintenance of whole-organism immunity across other cell types, most notably in immune cells. We observed no differences in gene expression in the *O. arbuscula* Immune Cell cluster between symbiotic and aposymbiotic tissues. The maintenance of immunity function across other cell types in *O. arbuscula* in and out of symbiosis provides nuance to the proposed immunity-nutrient tradeoff that symbiotic cnidarians must balance; that is, hosting algal symbionts within gastrodermal cells may not reduce whole-organism immunity. The immune system has evolved in different ways across metazoans. For example, in the three-spined stickleback *Gasterosteus aculeatus*, eight distinct immune cell states were identified with distinct abundances and transcriptomic profiles across populations, demonstrating specialized and rapid evolution of compartmentalized immune regulation (67). Here, we observe that “microevolution” of the immune system in *O. arbuscula* has occurred in three transcriptomically distinct immune cell states, indicating diversified immune cell function that is in agreement with other single-cell studies in stony coral (47). “Microevolution” may additionally occur in specialized algal-hosting gastrodermal cells, allowing these gastrodermal cells to function as pseudo-immune cells without affecting organismal immunity in immune cells. Notably, because canonical immune cells are able to maintain constitutive immunity across symbiotic states, previous work in field-collected tropical corals showing differences in pathogen susceptibility by symbiotic state (23,25) may be due to other factors (e.g., capacity for heterotrophy, efficiency of host-algal nutrient exchange, competition for nitrogen between host and algae) and not due to baseline immune cell tradeoffs due to symbiosis (26). However, additional work focused on characterizing differences between cell type-specific immune regulation in obligate versus facultative cnidarians is necessary to validate this hypothesis. Our comparative analysis between *O. arbuscula* and *Xenia* sp. (48) algal-hosting and non-algal-hosting gastrodermal cells suggests that obligate and facultatively symbiotic cnidarians may maintain their algal symbionts through different regulatory pathways, such as through classical immune response pathways in *Xenia* algal-hosting cells compared to cellular stress response pathways in *O. arbuscula* algal-hosting cells, although further research is needed to explore this hypothesis.

Our single-cell atlas also provides insights into carbon and nitrogen cycling within the gastrodermal cells of symbiotic and aposymbiotic *O. arbuscula*. Notably, we find that the gastrodermal cells that suppress immunity to allow for symbiosis are the same cells that undergo active nutrient cycling, most likely to regulate algal cell proliferation (11,14,68). Previous work has proposed that increased nitrogen cycling and assimilation occurring in symbiotic host cells controls algal cell proliferation to maintain optimal nutrient exchange between host and symbiont, preventing parasitism of the symbiont on the host (11,55). In our data, the co-expression of sugar transport and nitrogen cycling genes within the same symbiotic cells in specific gastrodermal cell clusters supports recent work showing that the stability of cnidarian-algal relationships relies on glucose-dependent nitrogen competition between the host and symbiont (12). In this model, the metabolites and energy produced from host glycolysis of symbiont-derived glucose may be used by the host for amino acid synthesis, through which nitrogen species, such as ammonium, can be assimilated. Hosts can thus modulate the amount of ammonium translocated to the symbiont for proliferation (12). However, in the present study, gastrodermal cells from aposymbiotic tissue exhibited lower expression of sugar transport and nitrogen cycling genes, which is likely driven by symbiotic cnidarians obtaining glucose sugars from symbiotic algae, whereas aposymbiotic *O. arbuscula* rely on heterotrophic sources to meet their metabolic needs (9,69). Thus, metabolic processing of carbon and nitrogen are likely performed through distinct pathways in cnidarians that rely predominantly on heterotrophy (i.e., aposymbiotic corals) compared to those that rely predominantly on autotrophy (i.e., symbiotic corals) (55).

Finally, while we observed no differences in the composition or diversity of the bacterial communities between aposymbiotic and symbiotic *O. arbuscula*, microbial partners may still play an important role in nutrient cycling and possibly host immunity. Differences in bacterial composition by symbiotic state may not have been observed in our study because corals were housed in common garden conditions. Future experiments with increased sample sizes collected *in situ* coupled with metabolomics or metatranscriptomics of the bacterial communities would more fully capture the functional variation of bacterial communities between symbiotic states.

Our scRNAseq approach applied in a facultative coral provides a novel way to study the tradeoffs and cellular regulation associated with the cnidarian-algal symbiosis. Previous work using whole-organism experimental techniques suggested that immunity costs are associated with hosting algal symbionts (14,20). Here, we showcase how differences in cell states and gene expression can conflate patterns observed in whole-organism ‘omics studies and, importantly, facilitate cell type-specific immune regulation to allow for endosymbiosis. Our description of highly specialized cell state-specific immune system regulation in a basal metazoan indicates that the immune costs of symbiosis previously observed in whole organism studies might not be as costly as previously assumed. Future work should compare how compartmentalized immunity is modulated in different host-algal pairings in facultative versus obligate cnidarians and also determine how these cellular processes are shaped by changing environmental conditions.

## Acknowledgements

This research was supported by a National Science Foundation grant IOS-1937650 (to T.D.G. and S.W.D.). M,V.-I. was supported by an NSF Graduate Research Fellowship, NSF NRT DGE 1735087, and a Boston University Marine Program (BUMP) Warren McLeod Marine Fellowship. J.K.D.-A. was supported by a BUMP Warren McLeod Marine Fellowship. We thank Diana Burkart-Waco of 10X Genomics Support for her assistance with data processing. We also thank Boston University Microarray & Sequencing Resource Core Facility, the Boston University Single Cell Sequencing Core Facility, Boston University Shared Computing Cluster, and the Tufts University Core Facility.

## Data availability

Raw fastq files for 16S and single-cell RNA-seq data can be found on NCBI’s Short Read Archive Bioproject PRJNA1122932. All data and code for analyses can be found at https://github.com/mariaingersoll/Oculina_sc_manuscript.git.

## Competing interests

The authors declare no competing interests.

## Notes

### Competing Interest Statement

The authors have declared no competing interest.

### Summary of Updates

1)We have performed a thorough bioinformatic comparison of mapping efficiency and quality control metrics of our single-cell data to two new references (a previously published transcriptome of the host coral Oculina arbuscula and a de novo assembled transcriptome from our single cell data) and concluded that the original Astrangia poculata genome provided higher quality mapping, as confirmed by several metrics. 2)We have conducted additional analyses to more confidently identify the cells hosting algal symbionts by analyzing reads mapping to the algal transcriptome. We have added further analysis demonstrating shared cell trajectories of symbiotic O. arbuscula gastrodermal cells to support our conclusions of algal-hosting cell identity. 3)We have included comparative analyses of gastrodermal algal-hosting cells from a previously published paper in the soft coral Xenia sp using orthologous genes. 4)We have provided additional methodological details and quality control testing in the main manuscript as well as additional supplemental figures and datasets. 5)We have included supplemental materials highlighting more in-depth analysis of all cell clusters identified in the manuscript. 6)We toned down some of the language around immune system compartmentalization while simultaneously bolstering our findings with additional analyses. We changed the title of the manuscript to reflect these changes 7)We have published a protocols.io methodology (DOI will be available pending manuscript acceptance) detailing the sample preparation and cell isolation.

