## Supplemental Information for "Cell type-specific immune regulation under symbiosis in a facultatively symbiotic coral"

**Competing interests:** The authors declare no competing interests.

**Contents**

Supplementary Figures S1, S2, S3, S4, S5, S6, S7

Supplementary Tables 1, 2, 3

Supplementary Materials and Methods

**SUPPLEMENTARY FIGURES**

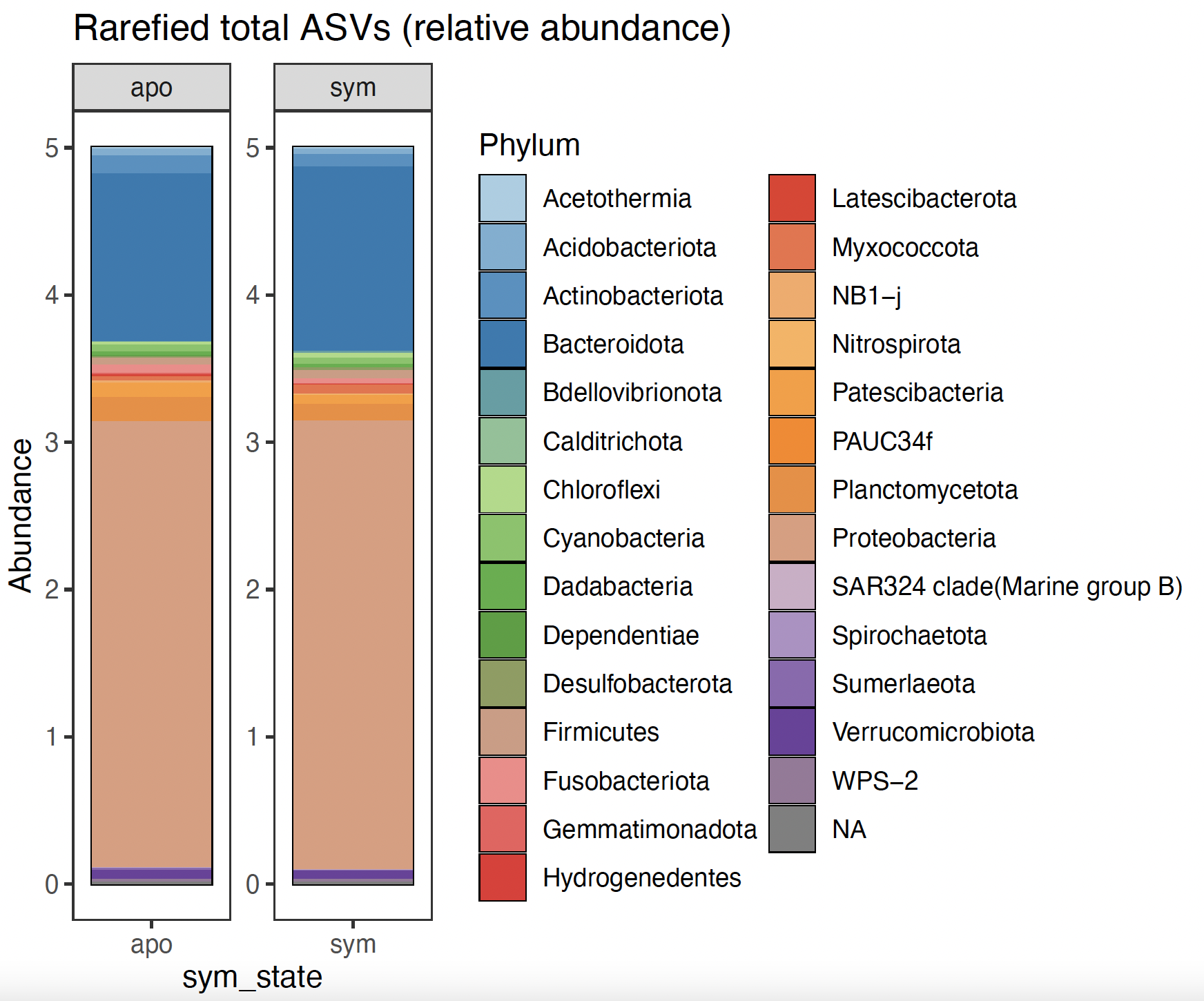

**Figure S1:** **Symbiotic and aposymbiotic *O. arbuscula* have similar bacterial communities.** Relative abundance of 16S ASVs from rarefied metabarcoding reads across different bacterial Phyla in symbiotic and aposymbiotic *O. arbuscula*.

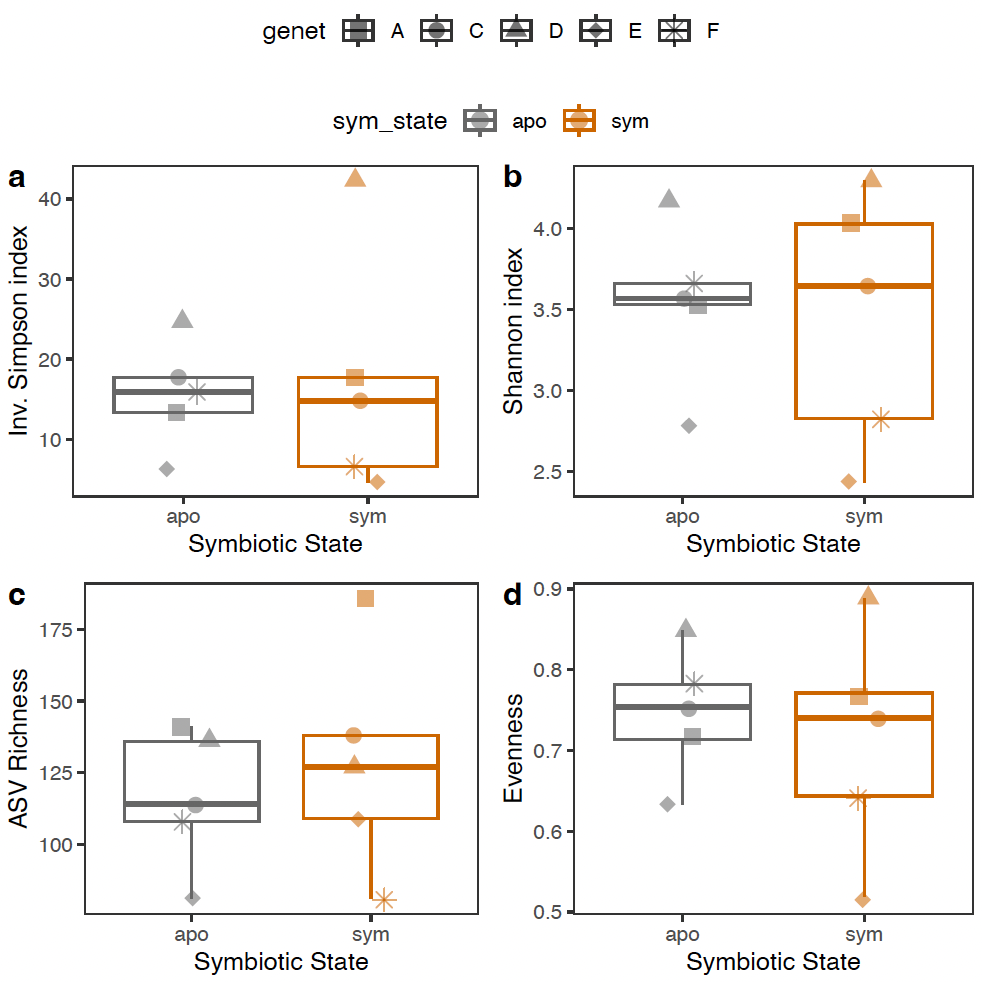

**Figure S2: No differences in bacterial diversity between symbiotic and aposymbiotic *O. arbuscula.*** 16S metabarcoding data showing alpha diversity (**a** Simpson Diversity; **b** Shannon Diversity, **c** Observed Species Richness from rarefied reads; **d** Evenness) between symbiotic and aposymbiotic *O. arbuscula*. Box plots display the median and the 1st and 3rd quartiles (hinges). Whiskers extend to the maximum and minimum points if they fall within 1.5 x the Interquartile Range (if no whiskers are drawn, points outside of the box are outliers). Each point is the alpha diversity metric of one individual (with the shapes representing genets).

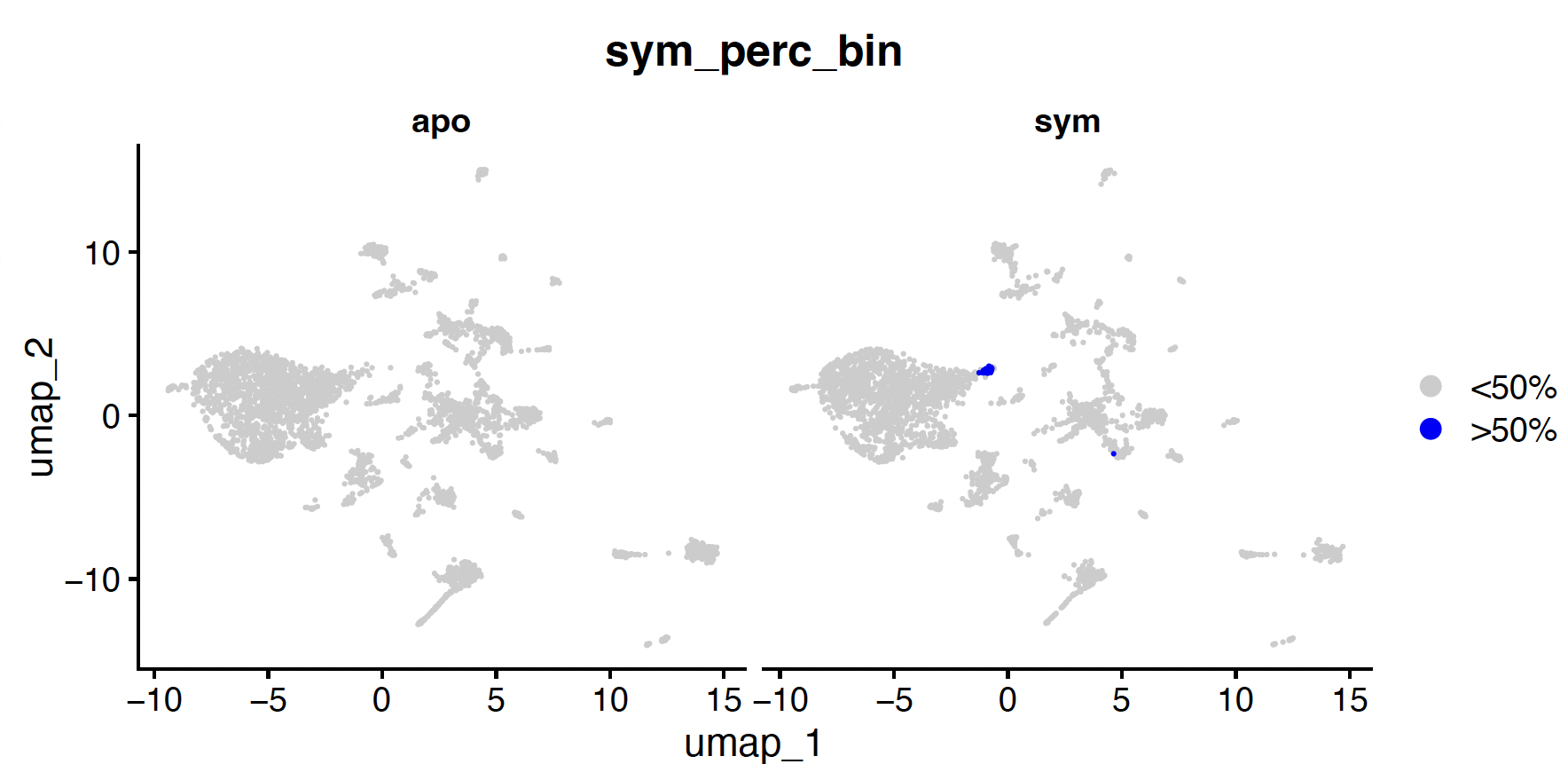

**Figure S3: Algal cells identified in Gastrodermis 2 cluster from symbiotic *O. arbuscula.*** Cells in which over 50% the reads corresponded to *B. psygmophilum* genes are highlighted in blue. 34 cells were identified, 33 of which were found in the Gastrodermis 2 cell cluster in the symbiotic sample.

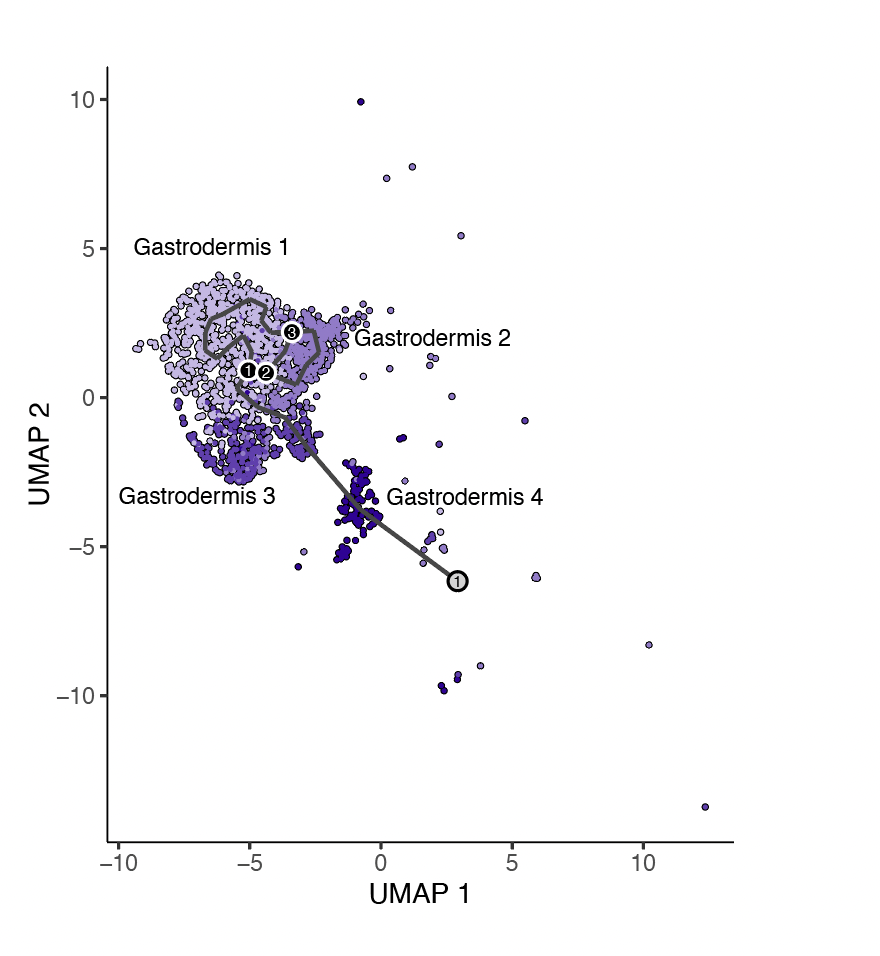

**Figure S4: Gastrodermis cells from aposymbiotic *O. arbuscula* have a single outcome.** UMAP projection of Gastrodermis 1, 2, 3, and 4 cells from aposymbiotic *O. arbuscula* overlayed with a graph of cell trajectories. The single cell outcome (fates) is denoted by a gray circle with black numbers. Branch nodes are denoted by black circles with white numbers.

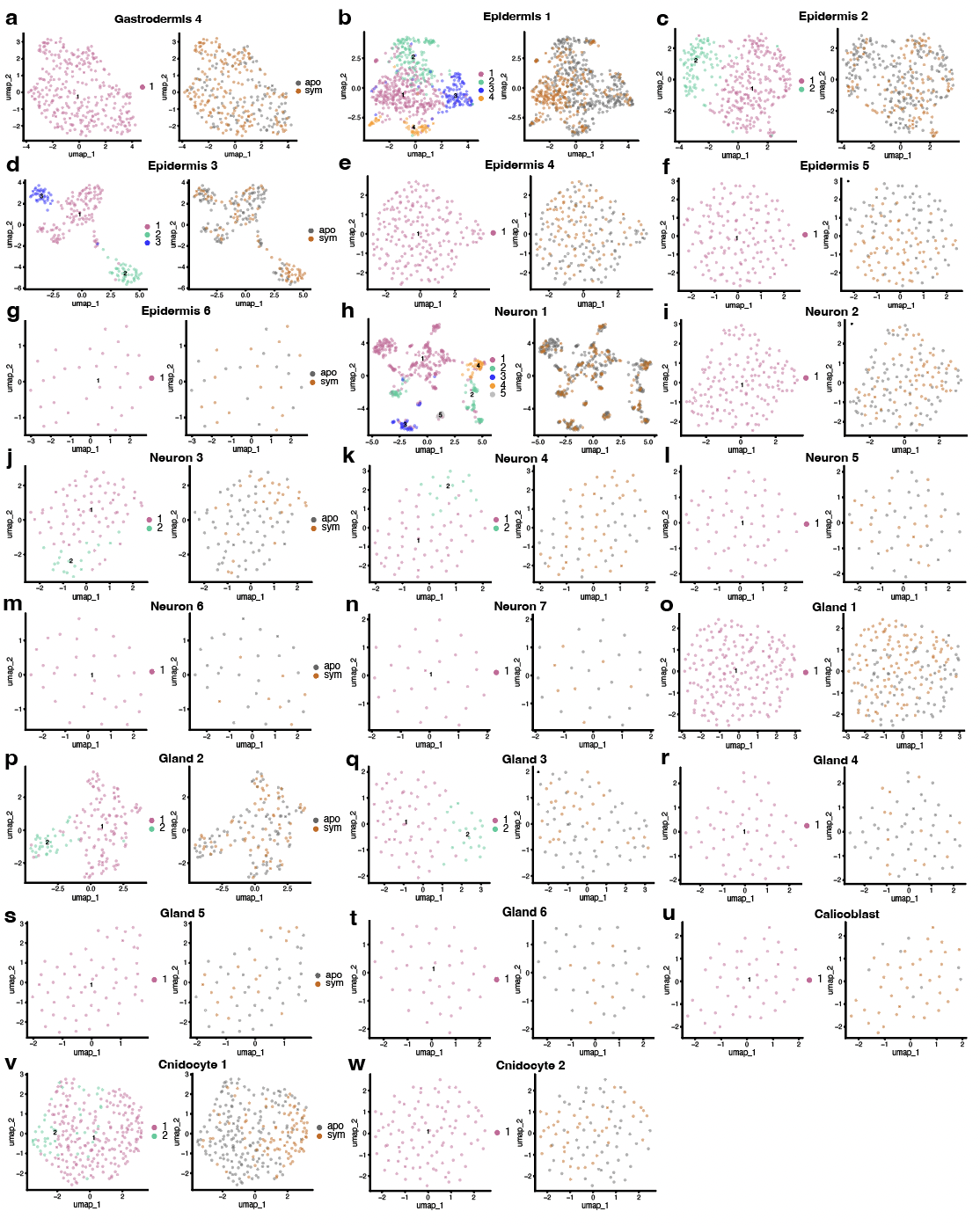

**Figure S5: Separation of cell clusters between subclusters and symbiotic states within different cell types.** UMAP projections of subsetted and reclustered cells from Gastrodermis 4 (**a**), Epidermis 1 (**b**), Epidermis 2 (**c**), Epidermis 3 (**d**), Epidermis 4 (**e**), Epidermis 5 (**f**), Epidermis 6 (**g**), Neuron 1 (**h**), Neuron 2 (**i**), Neuron 3 (**j**), Neuron 4 (**k**), Neuron 5 (**l**), Neuron 6 (**m**), Neuron 7 (**n**), Gland 1 (**o**), Gland 2 (**p**), Gland 3 (**q**), Gland 4 (**r**), Gland 5 (**s**), Gland 6 (**t**), Calicoblast (**u**), Cnidocyte 1 (**v**), Cnidocyte 2 (**w**). For each cell type, subclusters are color-coded in the left panel, and the symbiotic state of each cell is colored in the right panel.

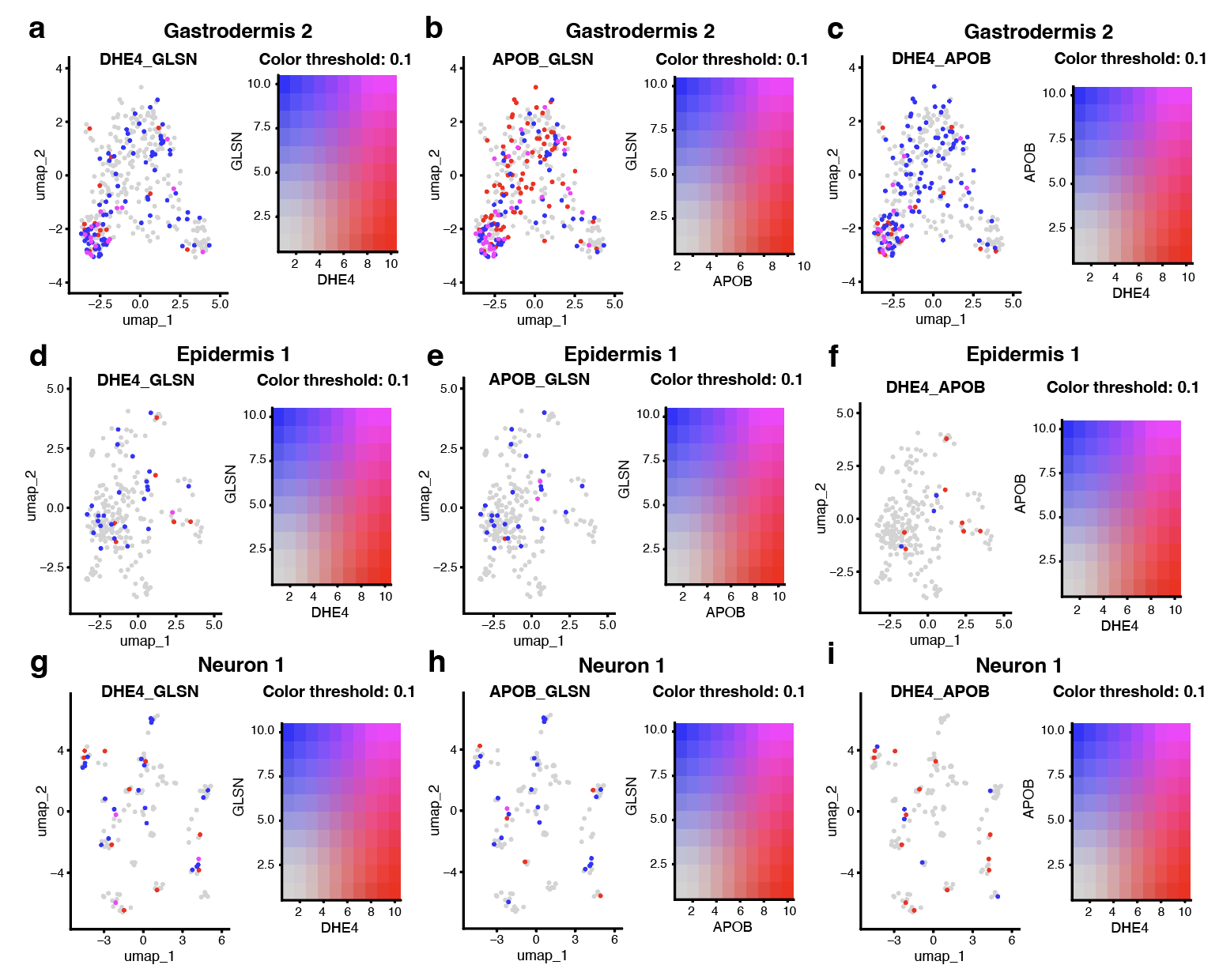

**Figure S6: Nutrient cycling genes of interest show little coexpression in Epidermis 1 and Neuron 1 cell clusters relative to Gastrodermis 1 and 2.** Coexpression plots of DHE4 and GLSN, APOB and GLSN, and DHE4 and APOB across Gastrodermis 2 cells (**a, b, c**), Epidermis 1 cells (**d, e, f**), and Neuron 1 cells (**g, h, i**) from symbiotic *O. arbuscula.* The coloration of each cell represents the per-cell mean expression value of each gene scaled to a maximum value of 10.

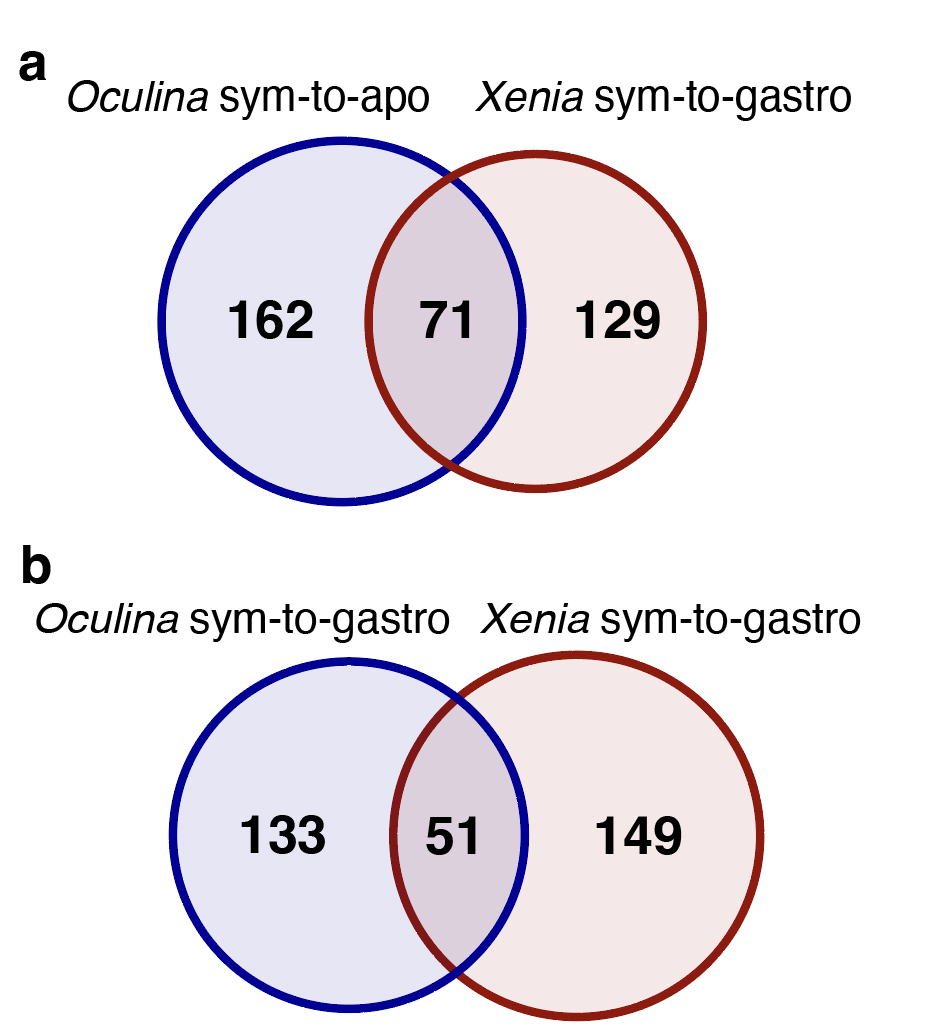

**Figure S7:** **Differentially expressed orthologs between *Xenia* and *O. arbuscula* gastrodermis and algal-hosting cells.** Overlap of differentially expressed orthologs (DEOs) between *O. arbuscula* Gastrodermis 1 and 2 cells from symbiotic and aposymbiotic tissue (*Oculina*-sym-to-apo) and *Xenia* algal-hosting and non-algal hosting gastrodermal cells (*Xenia*-sym-to-gastro) (a) and between *O. arbuscula* Gastrodermis 1 and 2 cells from symbiotic tissue and Gastrodermis 3 and 4 cells from symbiotic tissue (*Oculina*-sym-to-gastro) and *Xenia*-sym-to-gastro (b). *Xenia* data from (32).

**SUPPLEMENTARY TABLES**

**Table S1.** Biological Process GO terms involved in immunity that are differentially enriched between symbiotic and aposymbiotic *O. arbuscula* based on bulk analysis of scRNA-seq data.

| **Term** | **Name** | **p-adjusted** |
| --- | --- | --- |
| **GO:0043124** | **negative regulation of I-kappaB kinase/NF-kappaB signaling** | **0.06107341** |
| **GO:1901222** | **regulation of NIK/NF-kappaB signaling** | **0.08019282** |
| **GO:0043123** | **positive regulation of I-kappaB kinase/NF-kappaB signaling** | **0.09777471** |
| **GO:0002474** | **antigen processing and presentation of peptide antigen via MHC class I** | **0.003920727** |
| **GO:0050851** | **antigen receptor-mediated signaling pathway** | **0.07557286** |
| **GO:0050852** | **antigen receptor-mediated signaling pathway** | **0.07557286** |
| **GO:0002429**  **GO:0002768** | **immune response-regulating cell surface receptor signaling pathway** | **0.0003672095** |
| **GO:0002253**  **GO:0002757**  **GO:0002764** | **activation of immune response** | **0.008564266** |
| **GO:0050778** | **positive regulation of immune response** | **0.008848237** |
| **GO:0045087** | **innate immune response** | **0.07541165** |
| **GO:0002699** | **positive regulation of immune effector process** | **0.08770924** |
| **GO:0002684**  **GO:0050776** | **positive regulation of immune system process** | **0.09461819** |
| **GO:0002251**  **GO:0002385** | **organ or tissue specific immune response** | **0.09633996** |

**Table S2.** Resolutions used to define subclusters within each cell cluster of the full dataset.

| **Cluster** | **Resolution** |
| --- | --- |
| Gastrodermis 1 | 0.25 |
| Gastrodermis 2 | 0.25 |
| Gastrodermis 3 | 0.3 |
| Gastrodermis 4 | 0.3 |
| Epidermis 1 | 0.5 |
| Epidermis 2 | 0.3 |
| Epidermis 3 | 0.3 |
| Epidermis 4 | 0.3 |
| Epidermis 5 | 0.3 |
| Epidermis 6 | 0.3 |
| Neuron 1 | 0.3 |
| Neuron 2 | 0.3 |
| Neuron 3 | 0.3 |
| Neuron 4 | 0.3 |
| Neuron 5 | 0.5 |
| Neuron 6 | 0.5 |
| Neuron 7 | 0.5 |
| Gland 1 | 0.3 |
| Gland 2 | 0.3 |
| Gland 3 | 0.3 |
| Gland 4 | 0.3 |
| Gland 5 | 0.3 |
| Gland 6 | 0.3 |
| Calicoblast | 0.3 |
| Cnidocyte 1 | 0.3 |
| Cnidocyte 2 | 0.3 |
| Immune Cell | 0.4 |

**Table S3.** Raw and filtered read counts for samples used in 16S profiling (sym=symbiotic and apo=aposymbiotic).

| **Sample** | **Sym State** | **Raw** | **Filtered** |
| --- | --- | --- | --- |
| A13 | Sym | 21357 | 20430 |
| A8 | Apo | 21478 | 20255 |
| F8 | Apo | 10073 | 9138 |
| F1 | Sym | 13139 | 12627 |
| D7 | Sym | 14345 | 13561 |
| D6 | Apo | 21389 | 20513 |
| E8 | Apo | 13400 | 12994 |
| E4 | Sym | 65019 | 63410 |
| C6 | Apo | 14825 | 14102 |
| C9 | Sym | 13305 | 11219 |

**SUPPLEMENTARY MATERIALS AND METHODS**

Detailed information on the sample preparation, bioinformatic analyses, and R packages necessary to complete the experiments outlined in the Materials and Methods section of the manuscript *Cell type-specific immune regulation under symbiosis in a facultatively symbiotic coral*.

### **Coral husbandry and manipulation of symbiotic state**

Symbiotic (in symbiosis with *Brevioulum psygmophilum*) (1) colonies from seven genetic backgrounds (genets A-G) of *Oculina arbuscula* were collected at Radio Island Jetty, North Carolina (34˚ 42.520’ N, 76˚ 40.796’ W) in May 2018 under NC Division of Marine Fisheries Permit #1627488. These colonies have been maintained at Boston University in common garden aquaria (25 °C (± 0.17, SD), 34.6 PSU (± 0.76, SD), pH 8.0 (± 0.08, SD)) since May, 2018. One subset of aposymbiotic *O. arbuscula* branches was generated via menthol bleaching in Spring 2021 (symbiont counts, 16S metabarcoding, proteomics) and a second subset was generated in Spring 2022 (scRNAseq). Briefly, branches were incubated in 0.58 mM menthol in seawater under common garden light conditions. The menthol solution was replaced every 24 h for a total of two weeks. After aposymbiotic status was confirmed by a lack of symbiont autofluorescence under fluorescence microscopy (Leica M165 FC), aposymbiotic branches were transferred back to common garden aquaria and fragments were maintained for at least 2 months of recovery prior to physiological and multiomic profiling.

**Tissue removal for symbiont cell quantification**

To measure symbiont cell densities hosted in branches of symbiotic and aposymbiotic *O. arbuscula*, branches of approximately 2.5 cm in length were fragmented from symbiotic (N=10) and aposymbiotic (N=10) samples from genets A, C, D, E, and F, with at least one sample per genet for each symbiotic state. Tissue was removed via airbrushing into 0.2 μM-filtered Artificial Seawater (Instant Ocean). Total volume was noted, the resulting tissue slurry was homogenized, and an aliquot was subsampled for symbiont quantification using a hemocytometer. Skeleton surface area was measured using an Einscan-SE scanner and MeshLab software and symbiont cells were normalized to surface area. Statistical differences between symbiotic and aposymbiotic *O. arbuscula* fragments were calculated using the Kruskal-Wallis rank sum test in Rstudio v4.3.0, as assumptions of normality were not met.

**Coral spectroscopic determinations**

The light absorption capacity of symbiotic and aposymbiotic *O. arbuscula* was compared using reflectance (R) and absorptance (A) (2–4). Coral reflectance (R), the fraction of light reflected by the sample, was measured between 400-750 nm in intact coral fragments using a miniature spectroradiometer (Flame-T-UV-Vis, Ocean Optics Inc.). Briefly, samples were placed in a black container filled with filtered seawater and illuminated with homogeneous diffuse light by positioning a semi-sphere with an internal reflecting coating (barium oxide BaO) above the sample, and a ring of LEDs and halogen lamps directed upwards into the reflective coating. The reflected light was collected by a 2 mm diameter fiber-optic placed 1 cm above the surface of the sample at an angle of 45°. Reflectance was expressed as the ratio of the measurement from the tissue surface relative to the reflectance of a coral reference (a bleached *O. arbuscula* skeleton cleaned with commercial Hydrochloric Acid HCl). The coral absorptance (A), which describes the fraction of incident light absorbed by the coral tissue, was calculated from the reflectance spectra as A = 1 – R (2,4). The absorptance peak of chlorophyll a (Chl a) at 675 nm was calculated as A_675_ = 1 – R_675_, assuming that transmission through the skeleton of the samples is negligible.

**Microbiome profiling**

To identify bacterial communities associated with symbiotic and aposymbiotic fragments, one symbiotic and one aposymbiotic fragment from genotypes A, C, D, E, and F were flash frozen, and one polyp from each fragment was preserved in ethanol for 16S metabarcoding (N=10). Metabarcoding libraries were generated using a series of PCR amplifications for the V4/V5 region of the bacterial 16S rRNA gene (5,6) as follows: 95°C for 40 sec, 58°C for 120 sec, and 72°C for 60 sec for 32 cycles, with a final elongation step of 72°C for 5 min. PCR products were purified using GeneJET PCR Purification kits (ThermoFisher) and eluted in 30 µl. Each PCR product was barcoded via five PCR cycles and visualized on a 1% agarose gel to assess relative concentrations. Five negative controls using water were prepared and later used to remove contaminating sequences. Samples were pooled in equal concentrations, gel extracted, and submitted for paired-end 250 bp sequencing on an Illumina Miseq at Tufts University Core Facility.

16S primers were removed from raw reads using cutadapt (7). DADA2 v1.28.0 (8), quality filtering was conducted, and 1,232 sequence variants (ASVs) were inferred. Taxonomy was assigned at 100% sequence identity using the *Silva* v. 138.1 database (9). ASVs matching mitochondria, chloroplasts, or non-bacterial kingdoms were removed (83 ASVs removed) and 20 ASVs were removed based on negative controls as contaminants (decontam v1.2.0; (10)). Cleaned counts were rarefied to 8,298 using vegan v2.6-4 (11) and trimmed using MCMC.OTU v1.0.10 (12) to remove ASVs in less than 0.02% of counts, resulting in 661 ASVs across samples. The composition of bacterial communities across symbiotic state were compared via alpha diversity (Shannon index, Simpson’s index, ASV richness, and evenness) using phyloseq v1.44.0 (function *estimate_richness* (13)) and beta diversity was assessed using a PCoA on Bray–Curtis dissimilarity with a PERMANOVA in the vegan package (v2.6-4 (11)). Alpha diversity metrics were compared using linear mixed effects models ((package lme4; v1.1-35.3 (14)) with symbiotic state as the predictor and a random effect of coral genotype. The effect of symbiotic state on beta diversity was assessed using the function *betadisper* (vegan package; v2.6-4 (11)). DESeq2 v1.40.2 (15) then explored differentially abundant ASVs to ensure that subtle differences in specific taxa were not overlooked.

**Proteomic profiling**

Mass spectroscopy (MS) was used to identify differentially enriched proteins from total protein isolated from fragments of five symbiotic genets (Ax2, C, E, and F) and three aposymbiotic genets (C, E, and F). As described previously (16), each fragment (approximately 1 cm x 1 cm) was washed in PBS then crushed and incubated in 1X AT Lysis buffer with proteinase inhibitors (10 mM HEPES pH 7.9, 1 mM EDTA, I  mM EGTA, 20% (w/v) glycerol, 1% w/v Triton X-100, 20 mM NaF, 1 mM Na_4_P_2_O_7_·10H_2_O, 1 mM dithiothreitol, 1 mM phenylmethylsulfonyl fluoride, 1 μg/ml leupeptin, 1 μg/ml pepstatin A, 10 μg/ml aprotinin) for 1 h at 4 ℃ on an orbital shaker with occasional vortexing. The lysate was clarified by centrifugation at 13,000 rpm for 15 min at 4 ℃, and the supernatant was stored at -80 ℃ prior to analysis by MS.

For MS, tryptic peptide mixtures were analyzed by nano-scale high-performance liquid chromatography (Proxeon EASY-Nano system, Thermo Fisher Scientific) coupled with online nanoelectrospray ionization tandem MS (Q-Exactive HF-X mass spectrometer; Thermo Fisher Scientific). Briefly, samples were loaded into the system with aqueous 0.1% (v/v) formic acid via a trap column (75 μm i.d. × 2 cm, Acclaim PepMap100 C18 3 μm, 100 Å, Thermo Fisher Scientific) and peptides were resolved over an Easy-Spray analytical column (50 cm × 75 μm ID, PepMap RSLC C18, Thermo Fisher Scientific) by an increasingly mobile phase B. comprising 2% acetonitrile and 0.1% formic acid, while organic phase B consisted of 80% acetonitrile and 0.1% formic acid. Reverse phase separation was performed over 120 min at a flow rate of 300 nl/min. Eluted peptides were ionized directly into the mass spectrometer using a nanospray ion source. The mass spectrometer was operated in positive ion mode with a capillary temperature of 300 ℃ and a potential of 2,100 V applied to the frit. Tandem mass spectrometry (MS/MS) was performed using high-energy collision-induced dissociation, and 10 MS/MS data-dependent scans (45,000 resolution) were acquired in profile mode alongside each profile mode full-scan mass spectra (120,000 resolution), as previously described (17). For MS scans, the automatic gain control (AGC) was set at 1 × 10^6^ ions with a maximum fill time of 60 ms. MS/MS scans had an AGC of 3 × 10^4^, with a maximum injection time of 80 msec, activation time of 0.1 msec, and 33% normalized collision energy. To prevent repeated selection of peptides for MS/MS, a dynamic exclusion list was activated to exclude all fragmented ions for 60 sec.

For protein identification and analysis, data files (RAW format) were searched using the standard workflow of MaxQuant version 2.4 (<http://www.maxquant.org/>) under standard settings using *Astrangia poculata* genome (18). The parameters included two missed trypsin cleavage sites, fixed carbamidomethylation of cysteine, variable methionine oxidation, protein N-terminal acetylation, and phosphorylation of STY residues. For the first search, precursor ion tolerances were set at 20 ppm, and for the second search, they were set to 4.5 ppm. The MS/MS peaks were de-isotoped and searched using a 20-ppm mass tolerance. A stringent false discovery rate (FDR) threshold of 1% was used to filter candidate peptides and protein identifications. The searched intensity data were filtered, normalized, and clustered using Omics Notebook (19). Filtering was performed to remove any proteins not identified in at least 70% of samples, with 2,191 proteins passing the filter. After filtering, both datasets showed low levels of sparsity, and no missing value imputation was performed. The LIMMA R package was used for LOESS normalization and differential abundance analysis (20). Differential analysis of proteomic profiles between aposymbiotic and symbiotic samples was based on a moderated t-test (19). Proteins were considered differentially abundant if they had a Bonferroni adjusted P value < 0.1. Raw intensity counts were normalized using the *rlog* transformation function in DESeq2 (15) and visualized using a Principal Component Analysis (PCA) with the package vegan v2.6-4 (11). The effects of symbiotic state and genet were assessed using PERMANOVA with the adonis2 function in vegan v2.6-4 (11).

Predicted peptides of *O. arbuscula* were searched against the human proteome v.11.5 from the STRING v.11 database (21) with an e-value cut-off of 1x10^-5^. Protein–protein interactions of select differentially expressed proteins (FDR < 0.1) were retrieved from the STRING v.11 database (21). Interaction networks were visualized using Cytoscape v.3.7.2 (22).

**Single-cell RNA sequencing**

To create single-cell libraries, live cells from one symbiotic (F1) and one aposymbiotic (F8) branch were sampled from the same genet for single-cell isolation and 10X cDNA sequencing, using a previously verified protocol for coral cell isolation (23). As described on protocols.io at <http://dx.doi.org/10.17504/protocols.io.rm7vzkx72vx1/v1>, each fragment of approximately 1.5 cm in length was placed into 10 ml of ice-cold calcium-free artificial seawater (0.2-micron filter-sterilized). The fragment was then moved to 10 ml of ice-cold cell-isolation media (3.3x PBS, 2% heat-inactivated fetal bovine serum, 20 mM HEPES buffer in deionized H_2_O). Using RNase-free forceps, tissue was mechanically scraped from the skeleton until no visible tissue remained (<10 min). On ice, the cell-slurry was filtered through a 70-μm filter, then through two 40-μm filters. This process was repeated for each sample. 1 ml of filtered cell suspension was centrifuged at 4℃ for 10 min at 300 x g, and the pellet was resuspended in 100 μl of cell-isolation media with 0.2 U/μl Protector RNase Inhibitor. To maintain viability, cells were not sorted prior to processing. Cell isolation samples were analyzed by the Boston University Single Cell Sequencing Core Facility where cell counts and viability were quantified by 0.04% Trypan Blue staining and counting with a hemocytometer. Symbiotic branch F1 had a concentration of 2,625 cells/μl and a viability of 81.6%. Aposymbiotic branch F8 had a concentration of 3,312.5 cells/μl and a viability of 86.4%. These values are largely in line with other single cell preparation methods reported in the literature (viability threshold of 80%, *Nematostella vectensis* (24,25)).

cDNA for each sample was generated following the 10X Genomics Chromium Single Cell 3’v3 protocols, and quality was assessed via Bioanalyzer High Sensitivity DNA analysis. Samples were pooled in equimolar concentration, and the library was sequenced on an Illumina NextSeq 2000 (P3 100 kit) to obtain 50,000 reads per cell. 10X Genomics CellRanger (version 7.2.0 (26)) processed the sequencing reads, which were then aligned to concatenated genomes of *Astrangia poculata* (host) (18) and the algal symbiont *Breviolum psygmophilum* (27). Reads aligning confidently to the host and symbiont references were used to generate CellRanger output files. The *A. poculata* reference was used for mapping as there is currently no published genome for *O. arbuscula*, and alignment rates to a de novo assembled transcriptome from the 10X data generated here and to a previously published *O. arbuscula* transcriptome were of lower quality (Supplementary Dataset 1) (27–29). In addition, it is worth noting that species in the genera *Astrangia* and *Oculina* are phylogenetically related (28), and previous work has found that *O. arbuscula* populations from Radio Island, NC (source for this study) are more genetically similar to an *A. poculata* population from Woods Hole, MA (coral genome source) than they are to other, more sourthern *Oculina* populations (29). Default CellRanger cell calling algorithms were employed.

To create cell profiles for *O. arbuscula* and assign cell state identities, host reads from symbiotic and aposymbiotic samples were analyzed using Seurat (v.5.0.2) (30) in Rstudio (v.4.2.3). Genes expressed in fewer than three cells were discarded. Cells expressing fewer than 200 genes were removed, as were cells expressing greater than 3,000 genes (to remove possible doublets or multiplets). No duplicated barcodes were detected. Mitochondrial reads could not be removed, as the reference genomes/transcriptomes do not have annotated mitochondrial genomes. Datasets were first independently log-normalized, and the top 2000 genes that exhibited high cell-to-cell variation were identified, expression values of these genes were scaled, and a PCA was performed. To better identify cell clusters across both symbiotic states, the datasets were integrated using the Canonical Correlation Analysis (CCA) Integration method on the PCA reduction (nearest neighbors parameters). Clusters were identified using 30 dimensions and a resolution of 0.5 and were visualized using Uniform Manifold Approximation and Projection (UMAP). Marker genes were identified using FindAllMarkers (Wilcoxin Rank Sum Test; log_2_foldchange threshold of 0.5) and cell types were informed by gene annotations and marker gene comparisons to other cnidarian single-cell datasets (e.g., *S. pistallata* and *Xenia* sp.) (31,32). Expression patterns of top marker genes were assigned to cell clusters using violin plots, bubble plots, and visualization of gene expression in individual cells within the UMAP. 28 cell clusters across 7 cell types were identified in the UMAP. Lastly, to assess the broad transcriptomic differences between each cell cluster, the top 10 genes enriched in each cell cluster (using the average log_2_foldchange from FindAllMarkers) were identified.

To identify cells containing algal symbionts, we identified cells in which over 50% of the total reads corresponded to *B. psygmophilum* genes and then visualized these cells on the UMAP. These cells were then removed, as the low percentage of host reads confounded downstream analyses.

To determine how symbiosis alters enrichment of gene pathways at the whole-organism level, and how this differential enrichment is reflected in specific cell types, we performed Mann-Whitney U Gene Ontology (GO) term enrichment analysis (33). Significantly enriched GO terms within the ‘Biological Process’ (BP) GO division were defined with a false-discovery rate (FDR) of <0.1. Significantly over- and under-represented GO terms relating explicitly to immunity, antigen signaling, or the NF-κB pathway were identified (**Table S1**). Cells in which over 7% of the total reads corresponded to genes annotated with the GO terms of interest were visualized on the UMAP of all cells from symbiotic and aposymbiotic *O. arbuscula* samples.

To compare expression profiles of specific cell states between symbiotic and aposymbiotic samples, each cell cluster was independently reclustered, and the effect of symbiotic state on gene expression in each cell state was analyzed. First, the cell cluster of interest (*e.g.,* Immune Cell, Gastrodermis 1) was subsetted from the full dataset, and new variable features were identified. Data were then rescaled, and dimensionality reduction was re-run (using the first 30 dimensions). Subclusters were identified with resolutions between 0.25-0.5 (**Table** **S2**). The distribution of cells across the subclusters according to the symbiotic state was visualized using UMAPs. Differentially expressed genes between symbiotic states in all cell clusters were identified using DESeq2 within the FindMarkers function. Additionally, marker genes for each subcluster were identified. To highlight differences in the transcriptomes of the Immune Cell states, the top 20 genes enriched in each Immune Cell subcluster were identified and plotted (unannotated genes were removed).

We used the Monocle 3 package (v.1.3.7 (34)) to determine cell trajectories and outcomes for all gastrodermal cells from symbiotic and aposymbiotic samples. We plotted the UMAPs of the symbiotic and aposymbiotic subsets of Gastrodermis clusters 1-4 and overlaid the trajectory graph, noting nodes and outcomes.

To compare expression of genes involved in sugar transport and nutrient cycling in gastrodermal cells from symbiotic and aposymbiotic samples, the expression of sugar transport and nitrogen cycling genes in the reclustered Gastrodermis 1 and Gastrodermis 2 cells was analyzed. First, normalized expression values of genes involved in nitrogen cycling/symbiont density control (Glutamate Dehydrogenase DHE4 and Glutamate Synthase [NADH] GLSN) and sugar transport (Apolipophorin APOB) – identified in previous whole-organism RNA-seq studies (27) – were compared across all cell clusters using violin plots (function VlnPlt in the Seurat package (30)).. FeaturePlot (30) was then used to visualize the co-expression of these genes across the subclusters of Gastrodermis 1 and Gastrodermis 2 cells from the symbiotic sample (scaling the per-cell mean expression value of each gene to a maximum value of 10). To test whether the coexpression observed in the symbiotic gastrodermal cells is ubiquitous across other cell types, we also visualized coexpression of DHE4, APOB, and GLSN across Epidermis 1 and Neuron 1 from symbiotic tissue (scaling per-cell mean expression to a maximum value of 10).
